# Methylothon: a versatile course-based high school research experience in microbiology and bioinformatics-- with pink bacteria

**DOI:** 10.1101/2021.09.08.459370

**Authors:** Peyton A. Jones, David Frischer, Shannon Mueller, Thomas Le, Anya Schwanes, Alekhya Govindaraju, Katie Shalvarjian, Jean-Baptiste Leducq, Christopher J. Marx, N. Cecilia Martinez-Gomez, Jessica A. Lee

**Affiliations:** Integrative Biology Program, Harvard University, Cambridge, MA; Abraham Lincoln High School, San Francisco, CA; Berkeley High School, Berkeley, CA; Molecular and Cell Biology Program, University of California-Berkeley, Berkeley, CA; Galileo High School, San Francisco, CA; Plant and Microbial Biology Program, University of California-Berkeley, Berkeley, CA; Department of Biological Sciences, University of Idaho, Moscow, ID; Department of Plant and Microbial Biology, University of California-Berkeley, Berkeley, CA; Laboratory for Research in Complex Systems, San Francisco, CA; Space Biosciences Research Branch, NASA Ames Research Center, Moffett Field, CA

**Keywords:** project-based learning, course-based research experience, microbial ecology, bioinformatics, methylotrophy, community science

## Abstract

Methylothon is an inquiry-based high school learning module in microbial ecology, molecular biology, and bioinformatics that centers around pink-pigmented plant-associated methylotrophic bacteria. Here we present an overview of the module’s learning goals, describe course resources (available for public use on http://methylothon.com), and relate lessons learned from adapting Methylothon for remote learning during the pandemic in spring of 2021. The original in-person version of the module allows students to isolate their own strains of methylotrophic bacteria from plants they sample from the environment, to identify these using PCR, sequencing, and phylogenetic analysis, and to contribute their strains to original research in a university lab. The adapted version strengthens the focus on bioinformatics and increases its flexibility and accessibility by making the lab portion optional and adopting free web-based tools. Student feedback and graded assignments from Spring 2021 revealed that the lesson was especially effective at introducing the concepts of BLAST and phylogenetic trees, and that students valued and felt inspired by the opportunity to conduct hands-on work and to participate in community science.

## INTRODUCTION

Biological science education has embraced inquiry-based learning at a national level in recent years (1–3). This is due to its potential to improve enthusiasm for and learning retention in sciences, technology, engineering, and math (STEM), and to increase confidence and generate learning gains in important skills, especially for students from underrepresented minorities (URM) or students who are the least academically prepared (4, 5). In particular, Course-based Undergraduate Research Experiences (CUREs), in which a whole class engages in a research question or problem that is of interest to the scientific community, have the potential to achieve these advances (6–8). Notably, much of the published work on inquiry-based learning in the classroom has focused on the college students. High school students’ exposure to authentic research is often limited to summer programs, as high school classes lack the resources that many universities have to engage students in ongoing research, and may have stricter schedules that limit teachers’ freedom to introduce time-consuming experiments. Yet inquiry-based learning experiences have the potential to generate the same benefits in high school students as in undergraduates. Moreover, direct engagement in STEM activities and interaction with active STEM practitioners in a mentorship setting can help to widen students’ understanding of who can be a scientist (9) and create further bridges between classrooms and STEM opportunities in the community (10).

### Intended audience

Here we present Methylothon, a high school learning module in microbial ecology and evolution, structured similarly to a CURE but designed for 11th- and 12th-graders, and explicitly made flexible to suit a wide range of life sciences students. Methylothon was originally designed as a “nose-to-tail” microbial ecology module for high school biotechnology or biology classes, in which students would begin by isolating organisms from their local environment, proceed through PCR and sequencing for identification, and finish by learning phylogenetic tools to place their isolates on the tree of life, similar in principle to SEA-PHAGES or PARE (11, 12). The module includes a community science component, as isolates can subsequently be used in original research by collaborating researchers (our co-authors). Optional final projects would allow students to delve more deeply into studying microbial diversity in the context of biotech, or discuss difficult questions in research design. Methylothon thus provides a scaffold for building diverse skills and experiences, including hypothesis formulation, field work, plant identification, laboratory techniques in microbiology and molecular biology, bioinformatics, biological diversity/phylogenetics, science communication, and literature review. It touches on several of the core concepts for biological literacy outline in the AAAS 2011 Vision & Change report (Evolution; Information Flow; and Systems) and has potential to cover all six of the report’s listed core competencies (process of science; quantitative reasoning; modeling and simulation; interdisciplinarity; communication; understanding the relationship between science and society) (1), in addition to incorporating 17 concepts and skills from the American Society for Microbiology Curriculum Guidelines for Undergraduate Teaching (13) (Appendix 1).

Methylothon originated as an extension of an NSF-funded research project involving several of the co-authors on this paper, focused on understanding the ecology and evolution of the plant-associated methylotrophs of the genera *Methylobacterium* and *Methylorubrum*. As the research project involved isolating novel methylotrophs from the environment and characterizing them in the lab, we recognized that we could share our science with students by involving them in isolating methylotrophs as well. In addition to being extensively studied in the context of microbial single-carbon metabolism (14, 15), pink-pigmented facultative methylotrophs make ideal model organisms for introducing students to microbiology for a variety of reasons: they are ubiquitous in the environment, being found on a wide range of plant hosts (16, 17); they are very low risk to human health and to plant health; they are straightforward to isolate on selective media at room temperature; they have numerous biotechnological applications (17–21) and therefore appeal to students in biotech classes as well as those in more fundamental biology classes; and, finally, their pink pigmentation makes them easy to identify on agar plates.

Redesigning the module in 2020-2021 to meet the needs of remote instruction during the COVID-19 pandemic unfortunately required breaking some of the links between the students’ original data collection and data analysis, which are a valuable part of inquiry-based learning (22). However, it incurred the unexpected benefit of generating a course that is more versatile and more broadly accessible to classes without a molecular biology lab and/or with limited computing capacity. In this report we describe the remote-learning version of the lesson, but we have included in our online materials some resources for carrying out laboratory portions in class, including PCR protocols and recipes for culture media. Our goal is to enable teachers to choose their own approach depending on their learning goals and access to resources.

### Learning time

The full Methylothon learning module consists of 7 sessions. The timing of delivery can easily be varied according to the needs of the class, as evidenced by the contrasting approaches taken by our partnering high schools in 2021 (Table 1); our partner schools took between 1 and 3 weeks and some further extended with a final project at the end.

**Table 1.** The Methylothon curriculum can be taught over various timetables. Shown here are the lesson plans for two of our partner schools. In one class, the entire module was delivered in the course of a single week, as students met daily for lecture and additionally twice a week for lab sessions. Another class was able to devote only 2-3 periods per week to the module, so the full duration was three weeks.

| **1-Week Example Schedule** |  |  |
| --- | --- | --- |
| <u>Day</u> | <u>Period</u> | <u>Lesson</u> |
| 1 | morning | Guest Lecture; begin Leaf Press Lab |
| 2 | morning | Intro to Methylobacterium Lecture |
| 2 | afternoon | Tree of Life and Phylogenetics Lecture |
| 3 | morning | Bacterial Identification Lab (asynchronous) |
| 4 | morning | DNA Sequencing and Multiple Sequence Alignment Lecture; Bioinformatics Lab Walkthrough |
| 4 | afternoon | Bioinformatics Lab |
| 5 | morning | Finish Bioinformatics Lab; assign Final Project; conclude Leaf Press Lab |

| **3-Week Example Schedule** |  |  |
| --- | --- | --- |
| <u>Week</u> | <u>Day</u> | <u>Lesson</u> |
| 1 | 1 | Guest Lecture |
|  | 2 | Intro to Methylobacterium Lecture; begin Leaf Press Lab |
| 2 | 3 | Tree of Life and Phylogenetics Lecture |
|  | 4 | DNA Sequencing and Multiple Sequence Alignment Lecture; Bacterial Identification Lab (asynchronous) |
| 3 | 5 | Bioinformatics Lab Walkthrough |
|  | 6 | Bioinformatics Lab |
|  | 7 | Complete Bioinformatics Lab; assign Final Project; conclude Leaf Press Lab |

### Prerequisite student knowledge

Methylothon is designed for high school Juniors and Seniors; students should begin with some awareness of what microorganisms are, and understand what DNA is and be aware of the central dogma. Some of our partner classes already had experience with PCR, culturing microorganisms, or carrying out BLAST; for such classes, Methylothon can be a helpful review/synthesis activity. Alternatively, for classes where all the concepts covered are new, teachers may want to use Methylothon as a scaffold for just-in-time learning, and supplement with additional in-depth material between lessons to reinforce particular learning objectives. The structure of Methylothon offers substantial flexibility to the instructor to emphasize the learning objectives appropriate for their class; as an example, those marked with an asterisk on this list were topics of focus in some of the classes we taught but not all. The specific learning objectives listed here correlate with 17 of the key concepts and skills enumerated in the American Society of Microbiology Recommended Curriculum Guidelines for Undergraduate Microbiology Education (13); these are notated by their numbers in brackets for each learning objective, and described in Appendix 1.

### Learning objectives

By the end of Methylothon, students should be able to:

a. Describe the ubiquity and diversity of microbes in the environment [ASM 20, 27]
b. List 3 applications for microbes in biotechnology [ASM 23, 26]
c. Describe what methylotrophs are and what distinguishes them from other microbes [ASM 11, 12, 13]
d. Define what rare earth elements (lanthanides) are and identify the role they play in microbiological processes* [ASM 11, 12, 13]
e. Describe 3 methods used to identify microbes [ASM 34]
f. Explain how selective medium is used to culture methylotrophs [ASM 33]
g. Demonstrate standard methods for culturing microorganisms on agar culture plates and analyze the effects of methodological changes (the roles of temperature and moisture in colony development, etc.)* [ASM 33, 36, 37]
h. List the steps of PCR and explain the function of each ingredient [ASM 36]
i. Describe the role of the 16S rRNA gene in the study of evolutionary relationships [ASM 4, 5]
j. Run an NCBI BLAST analysis [ASM 34]
k. Describe what Multiple Sequence Alignment is and how it is used for understanding relatedness among microbes [ASM 5]
l. Analyze a phylogenetic tree and synthesize phylogeny with sample metadata to explore scientific questions [ASM 5]
m. Formulate and evaluate hypotheses, describe experimental procedures, and discuss sources of uncertainty for a microbial ecology experiment* [ASM 28, 29, 30, 38]

## PROCEDURE

The Methylothon learning module consists of lectures, homework assignments, and virtual labs, and, optionally, an in-person leaf press lab in schools that allow lab projects during remote learning (Fig. 1); all materials are available at http://methylothon.com. Also optional, but included in all our lessons taught in 2021, is an initial guest lecture by one of our team members who conducts microbiology research. Students are asked to prepare for the guest lecture by reading a blog post (23), profile, or even a scientific article written by the researcher (24) to prompt them to formulate questions ahead of time. Guest lectures feature details on current research in methylotrophy (e.g. the role of rare earth elements) or focus on microbial ecology and biotechnological applications more generally. Material covered in the guest lecture is not required for the remainder of the module; rather, the primary goal is as an entry event to engage students and spark interest in the topic, as well as providing an opportunity to interact with a practicing scientist.

**Figure 1.**
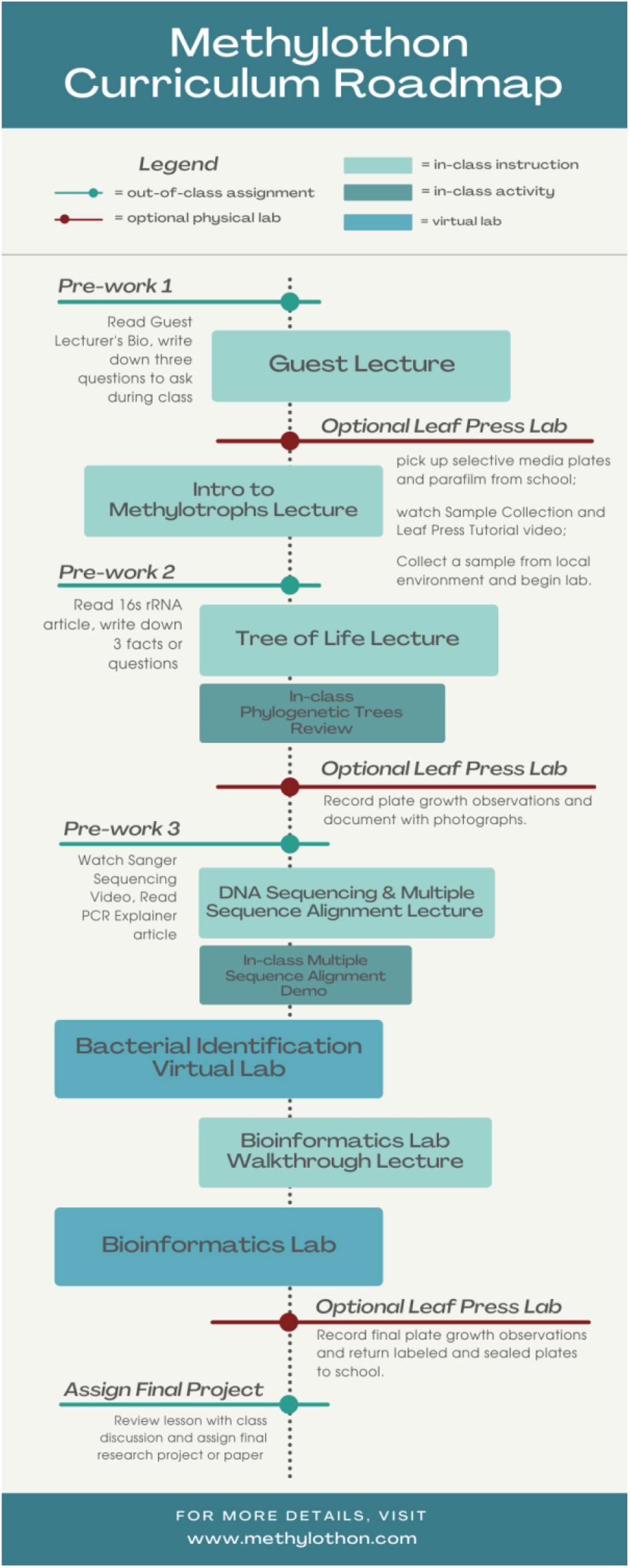
The components of Methylothon.

After the introductory guest lecture, the first official lesson of the module covers the motivation for working with methylotrophs and an overview of the process of sampling, isolating, and sequencing isolates, presented either as preparation for the leaf-press lab and an introduction to the community-science element of Methylothon (for classes that include the lab activity), or as background information on the origin of the sequences that students would soon analyze (for classes that omit it). A “how to make a leaf press” video is included in this lesson (available at http://methylothon.com) and is provided to students to watch again on their own as often as needed, providing a guide for techniques that might be new to students, such as the use of gloves, handling leaves without touching their surface, and wrapping parafilm around a culture plate.

For partner schools that include the leaf-press lab, it is assigned at this point as asynchronous work due by the following day. In the lab, students collect plant leaves from their neighborhood, press them gently onto two different selective media for culturing methylotrophs (recipe provided in Appendix 2), and incubate the plates, wrapped in parafilm, at room temperature in their homes. In 2021, our team provided teachers with culture plates, gloves, and parafilm, and they distributed them to their students with explicit instructions for storage and care. To explore the finding that some methylotrophs depend on rare earth elements for growth (as discussed in some guest lectures) (24), each student uses two plates: one containing lanthanum and one without. Incubation periods may last between 5 and 10 days, depending on the schedule of the class, though better results are obtained with >1 week incubation. At the end of each lesson, plates with bacterial colonies can be returned to one of our partner labs (in 2021, the Martinez-Gomez lab at UC Berkeley), where selected colonies can be further isolated and characterized. Students are required to submit photographs of plants and culture plates to an online repository (Fig. 2), and to record observations on dedicated online survey forms, including location and host plant identity (for this we encourage use of the augmented-reality app Seek by iNaturalist, https://www.inaturalist.org/pages/seek_app), and any growth observed on the culture plates by the end of the experiment.

**Figure 2.**
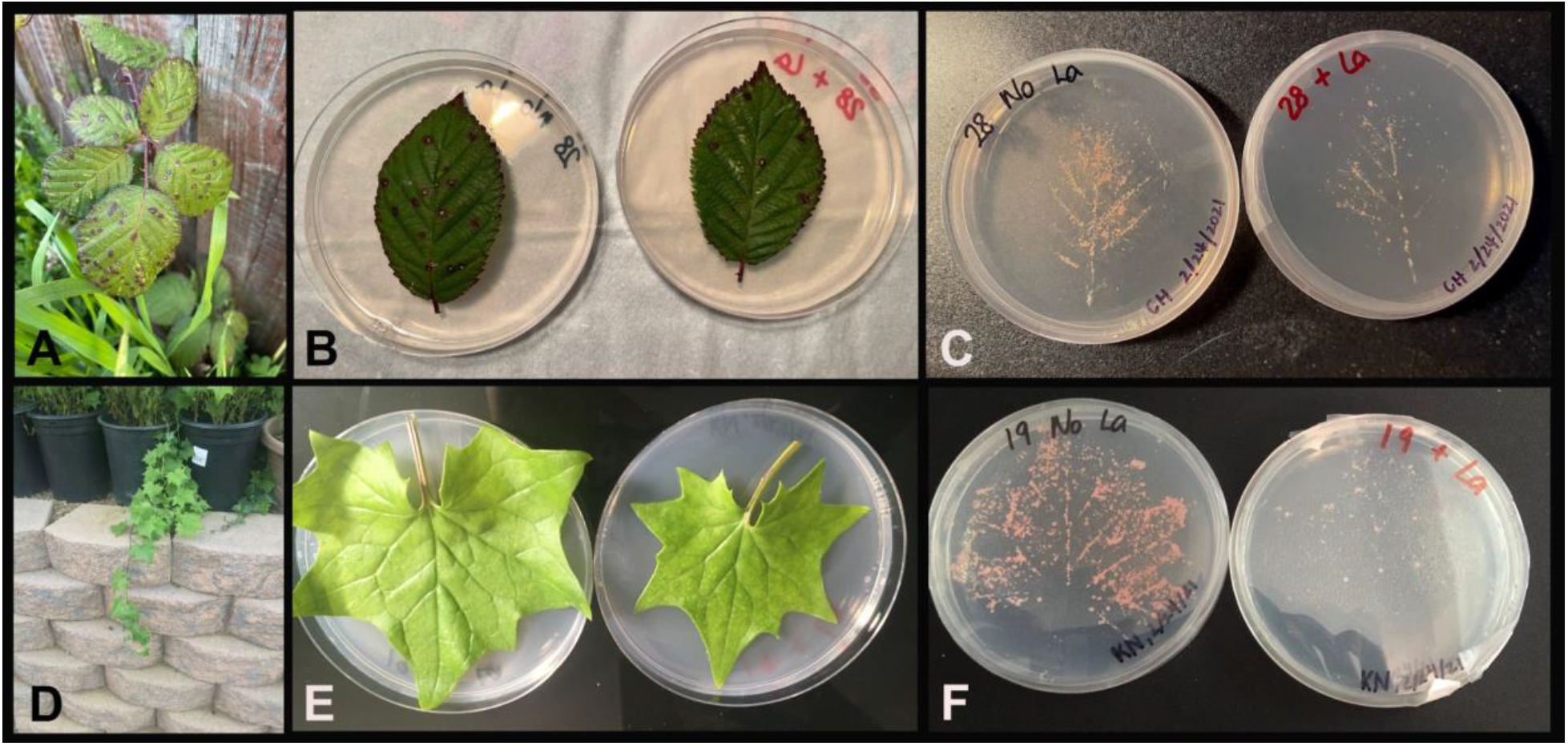
Selected photos of host plants and leaf press plates from Methylothon 2021. Both were collected in students’ backyards in San Francisco. A-C) Plant, leaf press, and colony growth from host identified as Brambles (likely Himalayan Blackberry, *Rubus armeniacus*). D-F) Plant, leaf press, and colony growth from host identified as Cape Ivy, *Delairea odorata*.

The second lecture covers the 16s rRNA gene and its role in microbial phylogeny. Students are assigned two brief articles from the microBEnet website as pre-reading (25, 26), and an interactive comment board, Google Jamboard (https://jamboard.google.com/), is used at the start of the lesson to review the homework. The lecture then discusses the three-domain system and the use of 16s rRNA as a standard for measuring the relatedness of bacteria and archaea. Students are led through a review of how to interpret relationships from phylogenetic trees, and use practice problems and breakout rooms to check understanding. The third lecture covers DNA sequencing and Multiple Sequence Alignment (MSA). Pre-assignments include short readings and video to review concepts on PCR and sequencing, which are reviewed in class using a Google Jamboard. The lecture covers interpretation of Sanger sequencing chromatogram and MSA, and includes an in-class exercise on the underlying concepts of sequence alignment.

In place of the PCR and sequencing that would be carried out lab during in-person instruction, students next complete a virtual lab exercise on bacterial identification produced by the Howard Hughes Medical Institute Biointeractive website (https://www.biointeractive.org/classroom-resources/bacterial-identification-virtual-lab), which allows students to simulate the procedure beginning with collecting a sample of a bacterial colony from a culture plate and ending with a “mini-BLAST’ of the sequenced DNA. This can be carried out as an asynchronous activity, or as synchronous group work; we provide worksheets for both settings to reinforce learning along each step of the lab.

In the final lecture, we walk through all the components of the Bioinformatics Lab that the students will ultimately complete on their own to understand the identity of a mystery methylotrophic organism. We introduce students to the NCBI BLAST online tool and its associated databases, and how to use free online tools to create sequence alignments and phylogenetic trees. To carry out the Bioinformatics Lab, students are given “mystery sequences”: FASTA files of 16S rRNA gene sequences from isolated methylotrophs, generated by previous classes. They are also provided with a file containing reference sequences of known methylotrophs and other bacteria for context. The students perform BLAST on their sequences for an initial identification, then perform MSA and construct a phylogenetic tree with the mystery sequences and reference sequences. Although each student has their own unique sequence, they work in small groups in online breakout rooms for peer support. Students complete a guided worksheet during the lab, and each ultimately uploads an image file of their phylogenetic tree to a class slideshow. The final worksheet questions require students to interpret their own and their classmates’ trees from the slideshow.

### Suggestions for determining student learning

The main formative assessments for Methylothon are the handouts completed by students during the two virtual labs, and a final project constitutes the summative assessment. Several other components of the module, including submission of photos and metadata for the leaf press lab, can be used for judging participation. During virtual labs conducted as group work, instructors frequently circulate among the breakout rooms as students work together, which provides additional opportunity for informal assessment of learning as well as evaluation of the module itself should changes be needed.

There is no single final project included in the Methylothon learning materials, but in 2021 each teacher developed their own in collaboration with the Methylothon team based on the specific learning goals for their class and the context within the rest of the curriculum, and examples are provided on the project website. As an example, students in one Biotechnology class that had carried out the Leaf Press Lab wrote a 2-page individual report summarizing the background, methods, results, and conclusions of the Methylothon module, including a section in which they researched literature to answer any question of their choice related to microbes in biotech (Appendix 3). In the International Baccalaureate (IB) Biology class, students worked in groups to write a report describing what they could and could not conclude from their Leaf Press Lab results, following IB International Assessment (IA) rubric guidelines, providing an opportunity for practice before their required IA report (Appendix 4). A subsequent assignment in that class involved writing 5-6-page individual essays that required them to combine their learning from Methylothon and from a previous Evolution unit to make an evidence-based argument and design an experiment demonstrating evolutionary principles. Similarly, another Biotechnology class incorporated principles from Methylothon with those from a unit on human ancestry in an assignment asking students to produce a report in the format of their choice (video, slideshow, essay, or graphic novel) on the process of analyzing and identifying biological samples using PCR, sequencing, and phylogenetics (Appendix 5). Pairing modules in this way emphasizes the universality of these methods and principles across the tree of life.

### Sample data

As mentioned above, student work in Methylothon includes both individual/group work that can be used for evaluating learning, and sample metadata and observations (collected via online survey forms) that can ultimately feed into the community-science component. Summaries of sample metadata from 2021 are shown in Fig. 3A (sample locations) and Appendix 6 (host plant species sampled), and examples of students’ detailed individual observations are provided in Table 2. Examples of students’ individual/group work for evaluation are provided in Appendix 7 (Bacterial Identification Lab) and Appendix 8 (Bioinformatics Lab).

**Figure 3.**
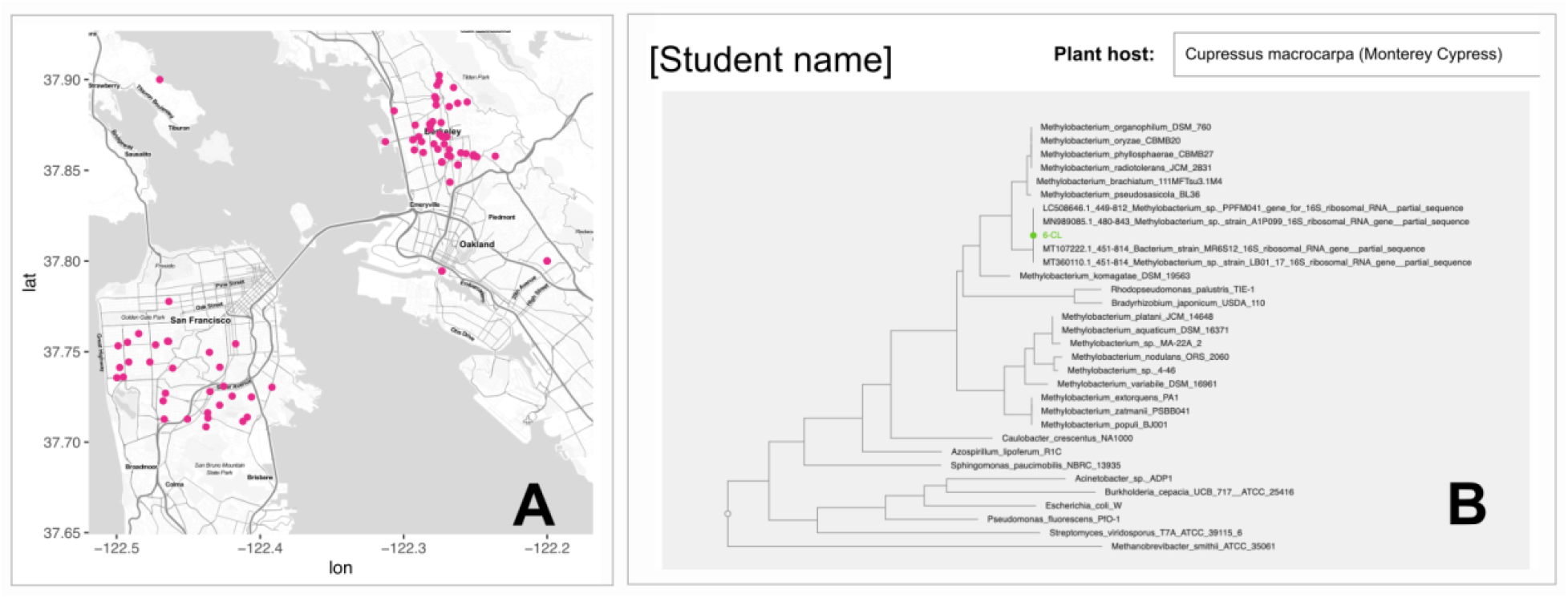
A) Sites sampled for plants by San Francisco and Berkeley high school students during 2021 Methylothon; each pink dot represents one sample site. B) Example of a student’s phylogenetic tree, in which the name of the student’s mystery sequence is in green text. Image taken from the class phylogenetic tree slide show; student’s name is omitted.

**Table 2.** Subset of the metadata accompanying the samples shown in Fig. 2, entered by students in the online form as part of the Leaf Press Lab. School name, sample date, and GPS coordinates are omitted.

| Sample code | Description of sample location | Other observations | Where in your house did you incubate your plate? | What is the general appearance of the colonies? |
| --- | --- | --- | --- | --- |
| CH 2242021 | The sample was taken in my backyard, which faces the ocean. My plant was surrounded by a grassy area with minimal flowers. Today was a relatively sunny day and the area where I sampled my leaves from gets a lot of sunlight. | The plant leaves have many thorns on the back and stems so when placing the leaves on the agar plate, the thorns pierced the agar. The rest of the process went smoothly. | On my desk in my bedroom. | The colonies are light pink, salmon-colored, and are crowded around the veins of the leaves. Overall, they are about the same size, but there are some that stick out more than others. On the plate without lanthanum, the colonies are hard to distinguish from each other because they are so packed together. |
| KN, 2/24/21 | The sample location was my backyard, which resides in a considerably chilly area of San Francisco, in the Excelsior district. The terrain is mountainous, and we grow chili peppers as well as numerous other things. There are many weeds and leaves of sorts. The amount of sunlight received within the area is decent, especially early afternoon. The sample grew from the ground and not in a pot. | Might have cracked the agar, but not entirely sure. Can clearly identify the outline of the plant on the agar, however. Process went smooth, in terms of sampling and identifying the plant. One of the leaves was a bit too big for the plate but somewhat worked. | I incubated my plate in my room, besides the window. It protects the plates from excessive sunlight but still allows some to shine in, therefore the agar would not dry out. The area is bit chilly and moist. | On the 19 No La plate, the colonies are all pink, some lighter in color than others. They take the form and shape of the leaf, and do not spread out too much. The colonies are very tightly packed and it is difficult to distinguish among them. On the other hand, the 19 + La plate has a few white colonies that are opaque, and the rest are light pink, much lighter than the colonies on the 19 No La plate. They are very spread out, compared to the 19 No La plate. |
| Sample code | Plant identification | Approximately how many colonies are there on the lanthanum side of the plate? | Approximately how many colonies are there on the NON-LANTHANUM side of the plate? |  |
| CH 2242021 | Brambles | About 100 colonies. | About 500 colonies |  |
| KN, 2/24/21 | Cape Ivy *Delairea odorata*<br>Genus: *Delairea* Family: Asteraceae<br>Classified within tribe Senecioneae Also known as German ivy in other parts of the world. | 30+ | 300+ |  |

### Safety issues

Methylothon carries relatively few safety risks, but instructors should be aware of a few key issues and their differences in the in-person and remote instruction cases. Typical field trip safety and liability issues apply for any class-based plant sampling trips. For in-person instruction, laboratory safety issues are the same as those typically associated with culturing Risk Group 1 microorganisms and molecular biology. Instructors should provide the necessary safety training according to the components of Methylothon they choose to implement; we refer readers to the ASM Guidelines for Biosafety in Teaching Laboratories (27), and particularly the “Addendum for biosafety considerations regarding at-home or DIY microbiology kits” for the remote version, with the caveat that there is no commercial kit provider to assume liability. Notably, we do not ask students at home to use high-proof ethanol, bunsen burners, or any of the other components of sterile technique that typically introduce laboratory hazards; moreover, once leaf press plates are wrapped in parafilm, there is no need for students to open the plates again, so the risk of contact with bacterial cultures, and of contamination of the plates, is low. One important and unavoidable hazard is the cycloheximide contained in the culture medium; we have found its inclusion to be necessary for inhibiting fungal growth sufficiently so that methylotrophic bacteria can be observed.

Cycloheximide carries some risk as a mutagenic and teratogenic agent. We therefore require that students wear gloves when handling culture plates. For remote instruction cases, students are required to return their culture plates to instructors, allowing for appropriate disposal in the lab.

Because sample metadata uploaded to methylothon.com are intended to be made publicly available alongside sequence data, student privacy should also be taken into account. Student names are collected to allow instructors to keep track of participation, but they are never published. If students choose to sample at their own homes, they are allowed to list GPS coordinates that are from the geographic vicinity but do not identify their home addresses.

## DISCUSSION

### Field testing

The in-person version of Methylothon was delivered two consecutive years (fall semesters 2018, 2019) to an Advanced Biotechnology class (seniors) at Abraham Lincoln High School (ALHS) in San Francisco, CA. During the COVID-19 pandemic in the 2020-2021 academic year, public high schools in the San Francisco Bay Area moved instruction online. We therefore modified Methylothon for remote learning, and at the same time took advantage of the online setting to expand the program’s reach to new schools. In addition to one section at ALHS, our spring 2021 partner schools included three sections of an IB Biology class (seniors) at Berkeley High School in Berkeley, CA, and four sections of a Biotechnology class (juniors and seniors) at Galileo High School in San Francisco, CA. Together, this entailed approximately 220 students and 55 hours of synchronous class time, with classes varying substantially in terms of learning goals and student experience level. COVID-related institutional policies also varied, such as the balance of synchronous versus asynchronous learning time, and the regulation enforced by some schools that students not be required to download software or register for online user accounts for bioinformatics tools.

### Evidence of student learning

As mentioned above, each of the classes that carried out Methylothon in 2021 designed their own final project as a summative assessment. As one example, students in one of the Biotechnology classes (3-week schedule) were assessed at several points throughout the module; formative assessments were given during the sequence that focused on PCR and bioinformatics, while summative assessments at the end of the sequence included an analysis of their phylogenetic trees and a final “abstract” writing assignment (Appendix 3). More than 70% of students showed early mastery of the concepts related to microbial culturing, PCR, DNA sequencing and DNA databases. Fewer students demonstrated mastery of the bioinformatics core concepts, particularly to the use of BLAST and phylogenetic tree generation, with most difficulties relating to the purpose of intermediate steps of the analysis pipeline. This was partly attributed to the fact that it was the students’ first exposure to these concepts; future implementations might include additional background learning and/or practice opportunities as a supplement to the Methylothon materials.

The IB Biology class (1-week schedule) assigned a summative assessment in which lab teams of 4 to complete a collaborative essay synthesizing their work from the module, including a hypothesis they had formulated and tested during the leaf press lab. Students were evaluated on their ability to incorporate background research to develop their research question and hypothesis, summarize their designed procedure, analyze their data quantitatively and qualitatively, develop a conclusion supported by their evidence, and evaluate their experimental design for sources of error and limitations (Appendix 4). The area that proved most challenging was the analysis of non-procedural sources of error and limitations of the experimental designs. Furthermore, because students were given only 1-2 days to develop their initial hypothesis, many did not have time to find, read, and analyze relevant scientific journal articles, but rather developed hypotheses based on background knowledge and information from the unit. However, all teams demonstrated achievement of the core objectives of the week, with 100% of the 21 teams receiving a grade of a 3.5 or higher (out of 4 total possible points), and 16 of the teams receiving a 4. The same class completed an additional qualitative assessment asking students to design an experiment that incorporated their understanding of the lecture material and utilized MSA, PCR, and BLAST to demonstrate microbial evolution. Over 90% of students successfully utilized the information they retained Methylothon, and many students went above and beyond by using bacteria outside of the methylotrophs to write their essays. We recommend that classes designing summative assessments accompanying Methylothon consider incorporating experimental design in the assessment, to enable students to continue developing these essential skills.

In addition to the school-specific assessments described above, the Methylothon team carried out a brief exercise to gather uniform feedback from all the classes we interacted with. Between five and eight weeks following our final lesson, we sent an anonymous 2-question survey to our partner teachers to distribute to their classes. In the survey, we asked students to tell us what they remembered most from the lesson and what they considered to be the most valuable takeaway from the lesson.

We received 79 responses. We grouped the responses into 7 categories based on the main topic referenced, plus an “Other” category (Fig. 4). Students reported remembering and valuing concepts relating to phylogenetic trees most often, followed closely by BLAST. We are encouraged by this result — most students remembered the goals around which we had framed the lesson! Additionally, we saw several students connect BLAST and building phylogenetic trees in their answers. To a lesser extent, students remembered intermediate steps such as DNA sequencing and the use of databases to retrieve information.

**Figure 4.**
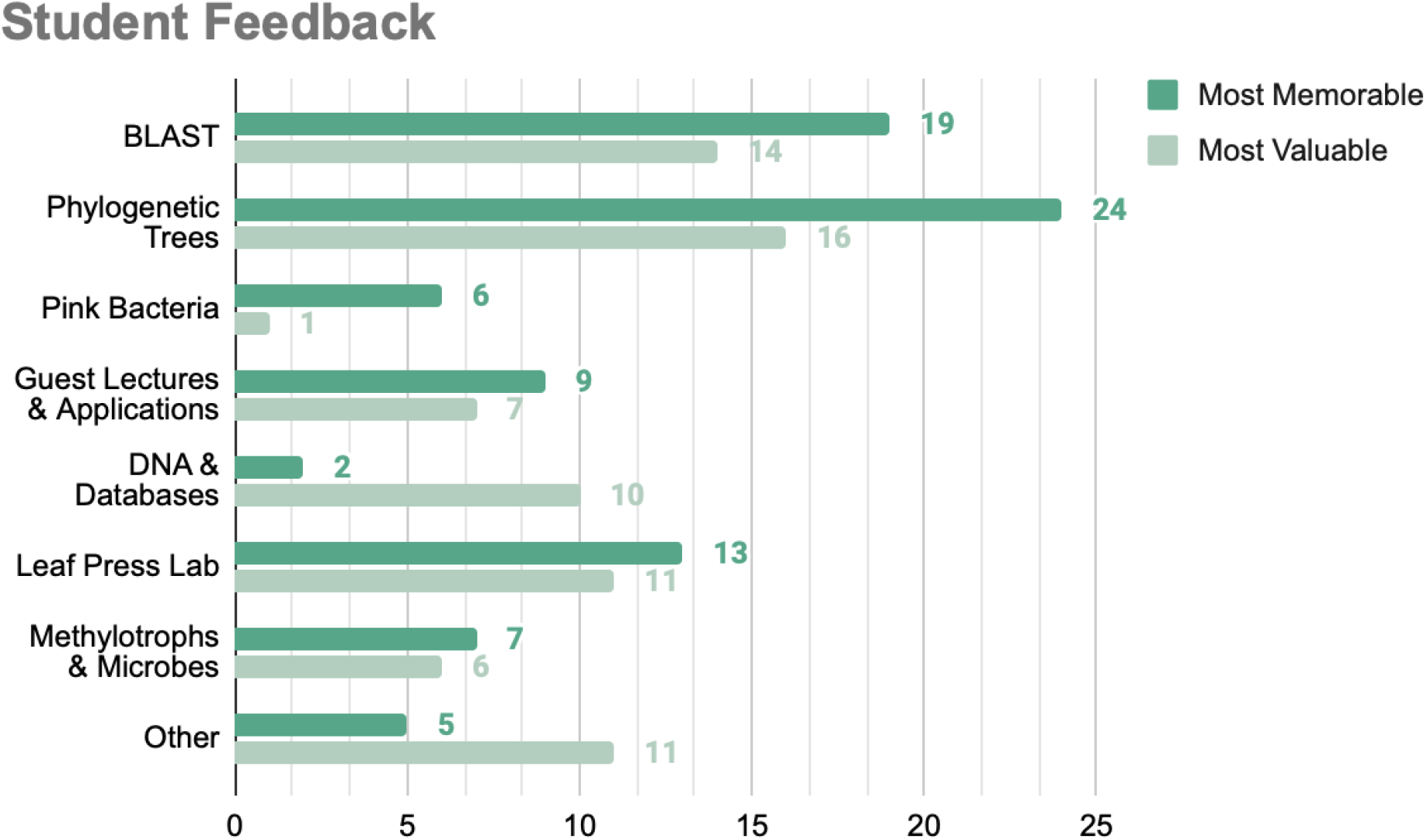
Topics mentioned in student responses collected from an anonymous survey that asked “What’s the one thing you remember most about the lesson?” and “What did you find most valuable about the lesson?”. Some responses incorporated multiple topics and were therefore counted more than once. Responses not fitting a predefined category were classified as “Other.”

In addition to the curriculum, students also recalled and valued other aspects of our lesson. Students who participated in the Leaf Press Lab (48 of the 79 respondents) responded positively to the chance to conduct some in-person lab work; 25% of those students reported that they most remembered the opportunity to do a non-virtual lab, and nearly 20% reported that the lab was what they found most valuable about Methylothon. Additionally, several of the students who had the opportunity to do the lab mentioned remembering that *Methylobacterium* were pink, reinforcing the importance of this phenotype in an educational model organism. In addition to the larger trends displayed in Fig. 2, some other facets highlighted by students as particularly helpful included the opportunity to speak to scientists about their educational background and research interests, and the chance to “experience what it is like to be a real scientist” (Box 1).

### Possible modifications

Teaching Methylothon to several different partner schools allowed us to identify several areas for improvement or modification. One of the major pivots we made in our later lessons was to provide DNA sequences in files with a .txt extension rather rather than .FASTA. A FASTA file is a plain-text file containing molecular sequence data in a specific format, and adding the .FASTA extension to the file can remind users of the format and help sequence analysis programs recognize the file. However, asking students — who did not have the necessary underlying knowledge of file formats, file extensions, and how to use text files that they could not open and view — to navigate opening and working with files with a .FASTA extension blurred the focus of the lesson, and created a host of computer errors to troubleshoot. The file type extension was not a main feature of the lesson, so after seeing multiple classes lose lesson time to FASTA issues, we chose to distribute the files with a .txt extension to our remaining classes.

Additionally, we found that many students were unsure of how to translate the concepts from our initial general lecture on phylogeny to their interpretation of the trees they generated, and many were unclear about the significance of the reference sequences. We therefore added in a quick walkthrough of what results to expect from the student-generated phylogenetic tree immediately after the students had shared their trees in the class slideshow and before they began interpretation. We found that providing the review before turning students loose to analyze their trees helped students approach interpreting their own results with more confidence.

Relatedly, we found that some classes would have done better with a lesson structure that was more focused on “just-in-time” learning. For classes in which Methylothon was stretched across multiple weeks, the elapsed time between initial lectures and hands-on practice in the bioinformatics lab was long enough that lecture material held less relevance for the students than we had hoped, and some concepts were forgotten before they were practiced. One partner teacher related that they would consider breaking up the bioinformatics lab next time so that students carried out individual steps immediately after the relevant lecture.

The execution of microbiology lab work in a remote-learning environment provided unique challenges. Teachers shouldered the burden of working with school administration to obtain approval for students to pick up and drop off materials. Yet the greatest challenge was in making the exercise a valuable scientific experience for the students, given that they would be unable to use molecular methods to identify their isolates. In one class, we not only discussed potential hypotheses and experimental design, but also the possible limitations and sources of error in the experimental setup (e.g. the limited number of plates available, the lack of controls, the short incubation time), and asked students to explore these limitations in their written report. For students who had done previous work with fast-growing organisms such as *Escherichia coli*, we also found it necessary to moderate their expectations by emphasizing how much more slowly methylotrophs grow. We recommend that classes attempting the leaf press lab allow students to observe their plates for at least a week before returning them to the teacher.

Finally, as the name Methylothon implies, this module was originally conceived, and can still be modified in the future, to incorporate an element of competition. For classes that are able to devote a longer time period to this module, students’ isolates could be cultured and “competed” in a variety of phenotypic assays (growth rate, temperature range, carbon substrate utilization, antibiotic resistance, tolerance to other stressors). A very wide range of phenotypic assays would provide a chance for every isolate to “win” at something, and the competition could be used to introduce concepts of ecological niche and evolutionary adaptation. Such assays might be carried out in schools if they have very well-equipped laboratories, or by partnering research laboratories (such as those of our co-authors) as part of the community-science aspect of the module, encouraging regular communication between the class and the research lab as students check in by email or video chat on the performance of their isolates.

## ACKNOWLEDGEMENTS

We are grateful to the members of the Martinez-Gomez Lab (UC Berkeley), particularly Nathan Good for assistance with handling methylotroph isolates in 2021. We thank the laboratory of José de la Torre (San Francisco State University) for assistance with the in-person version of Methylothon in 2018-2019, especially Brittany Baker for providing help with bioinformatics instruction. Methylothon is funded by a grant from the National Science Foundation Division of Environmental Biology, Dimensions of Biodiversity program (award 1831838), to C. J. Marx, N. C. Martinez-Gomez, and J. A. Lee.

## Box 1.

**Selected responses from the anonymous student survey.**

- “What I found most valuable about the lesson was honestly just being back in a lab oriented setting to a certain extent. Since COVID-19 has affected us in not doing many labs, I believe this lab was something that made me feel a bit more interested and motivated again.”
- “[What I valued most was] seeing the other student’s [phylogenetic tree] results to compare.”
- “[What I valued most was] getting to experience what it is like to be a real scientist.”
- “The lesson I found most valuable was being able to conduct the experiment, even if it was virtual, I think that really fun and engaging even over [Z]oom”
- “[What I valued most was] the chance to watch as pink bacteria grew on my petri dishes in the shape of the leaves, knowing that my data may potentially be used in a real scientific paper!”
- “I found the most valuable part of the lesson being learning about what is actually being researched right now in the field of biology.”
- “I found the introduction of BLAST the most valuable. It was my first time hearing of this term, and being taught how to utilize this program is a great skill I could carry with me in the future.”
- “The thing I found most valuable about the lesson was how easy it is to go outside and collect a sample to learn more about. “

