## Appendices 1-8 for "Methylothon: a versatile course-based high school research experience in microbiology and bioinformatics-- with pink bacteria"

**Appendix 1. Components of ASM Curriculum Guidelines (Merkel et al. 2012) covered in Methylothon.**

| | | |
|---|---|---|
| **Concepts and Statements** | | |
| | ***Evolution*** | |
| | 4 | The traditional concept of species is not readily applicable to microbes due to asexual reproduction and the frequent occurrence of horizontal gene transfer. |
| | 5 | The evolutionary relatedness of organisms is best reflected in phylogenetic trees |
| | ***Metabolic Pathways*** | |
| | 11 | Bacteria and Archaea exhibit extensive, and often unique, metabolic diversity |
| | 12 | The interactions of microorganisms among themselves and with their environment are determined by their metabolic abilities |
| | 13 | The survival and growth of any microorganism in a given environment depends on its metabolic characteristics. |
| | ***Microbial Systems*** | |
| | 20 | Microorganisms are ubiquitous and live in diverse and dynamic ecosystems. |
| | 23 | Microorganisms, cellular and viral, can interact with both human and nonhuman hosts in beneficial, neutral or detrimental ways. |
| | ***Impact of Microorganisms*** | |
| | 26 | Humans utilize and harness microorganisms and their products. |
| | 27 | Because the true diversity of microbial life is largely unknown, its effects and potential benefits have not been fully explored. |
| **Competencies and Skills** | | |
| | 28 | Ability to apply the process of science |
| | 29 | Ability to use quantitative reasoning |
| | 30 | Ability to communicate and collaborate with other disciplines |
| **Microbiology Laboratory Skills** | | |
| | 33 | Use pure culture and selective techniques to enrich for and isolate microorganisms. |
| | 34 | Use appropriate methods to identify microorganisms (media-based, molecular and serological). |
| | 36 | Use appropriate microbiological and molecular lab equipment and methods. |
| | 37 | Practice safe microbiology, using appropriate protective and emergency procedures. |
| | 38 | Document and report on experimental protocols, results and conclusions. |

#### Appendix 2. MP medium for Methylothon leaf press plates

**Yield:** 1000 ml (approximately 30-40 plates)
**Overview:** Make MP medium, a defined minimal medium optimized for methylotrophs, with agar. Just before pouring plates, add the appropriate carbon source, lanthanides, vitamins, and fungal inhibitors for leaf press plates.

**For MP (Modified PIPES) medium**
From Delaney et al., 2013. doi: 10.1371/journal.pone.0062957 (http://www.ncbi.nlm.nih.gov/pubmed/23646164)
Each batch of MP medium is composed from the following stock solutions. Full recipes and instructions for all stock solutions is in "Preparation," below.

| **Name** | **Stock concentration** | **Final concentration in MP medium** | **Volume to add for 1 L** |
|---|---|---|---|
| PIPES (10x) | 300 mM | 30 mM | 100.0 mL |
| P-Solution (100x) | 145 mM | 1.45 mM | 10.0 mL |
| $MgCl_2$ (4000x) | 2 M | 0.5 mM | 250 µL |
| $(NH_4)_2SO_4$ (250X) | 2 M | 8 mM | 4.0 mL |
| C7-metals (1000X) | 1.2 mM | 0.0012 mM | 1000 µL |
| $CaCl_2$ (100X) | 2 M | 0.02 mM | 10 µL |
| Bacto Agar | n/a | 15 g/L | 15 g |
| $ddH_2O$ | | | add to reach final volume |

**MP preparation**

| | |
|---|---|
| 1 | Prepare stock solutions as described in steps 2-8. If stock solutions are already available, skip to step 9. |
| 2 | To prepare PIPES stock solution, dissolve 90.711 g of PIPES free acid ($C_8H_8N_2O_6S_2$) in a final volume of 1 L of $dH_2O$.<br>Suggestion: add some KOH to ~700 mL of water before adding any PIPES. Measure out all the PIPES you will need. Add a small amount of PIPES at a time; the solution will turn milky-white until the PIPES dissolves. If all the PIPES dissolves, add more until it doesn't. Then add more KOH and repeat. Going back and forth between KOH and PIPES, ensure that all the PIPES dissolves; JUST BE CERTAIN YOU DON'T OVERDO THE KOH. After you've added and dissolved all the PIPES the pH should still be acidic. Carefully adjust the pH to 6.75 by adding KOH. Bring the volume up to 1 L and check pH again; adjust with more KOH if needed. |
| 3 | To prepare P stock solution, add 33.1 g of $K_2HPO_4$* $3H_2O$ and 25.9 g of $NaH_2PO_4$* $H_2O$ to 1 L of $dH_2O$. |
| 4 | To prepare the stock solution for $MgCl_2$ (magnesium chloride), add 81.32 g of $MgCl_2$* $6H_2O$ to 200 mL of $dH_2O$. |
| 5 | To prepare the stock solution for $(NH_4)_2SO_4$ (ammonium sulfate), add 52.9 g of $(NH_4)SO_4$ to 200 mL of $dH_2O$. |

| | |
|---|---|
| 6 | To prepare the stock solution for $CaCl_2$ (calcium chloride), add 58.8 g of $CaCl_2$ *$2H_2O$ to 200 mL of $dH_2O$. |
| 7 | To prepare the stock solution for C7 metals, prepare a container with 100 mL of $dH_2O$, and add each of the following. Add them in the order listed, being sure to dissolve each completely before adding the next.<br>1) 1341.1 mg of $Na_3C_6H_5O_7$ (sodium citrate)<br>2) 34.5 mg of $ZnSO_4$* $7H_2O$ (zinc sulfate heptahydrate)<br>3) 19.8 mg of $MnCl_2$* $4H_2O$ (manganese chloride tetrahydrate)<br>4) 500.4 mg of $FeSO_4$* $7H_2O$ (ferrous sulfate heptahydrate)<br>5) 247.1 mg of $(NH_4)_6Mo_7O_{24}$* $4H_2O$ (ammonium molybdate tetrahydrate)<br>6) 24.96 mg of $CuSO_4$* $5H_2O$ (copper sulfate pentahydrate)<br>7) 47.58 mg of $CoCl_2$* $6H_2O$ (cobalt chloride hexahydrate)<br>8) 10.88 mg of $Na_2WO_4$* $2H_2O$ (sodium tungstate dihydrate) |
| 8 | Autoclave all separate solutions. |
| 9 | To make the media, mix all the components together.<br>First, add the listed volumes from the stock solutions into a new bottle.<br>Then, in a volume of $ddH_2O$ sufficient to bring the medium to the final desired volume (calculate this based on the volumes of the supplements you will add after autoclaving - see below), melt the appropriate amount of Bacto Agar (15 g/L) using a stir bar and hot plate. DO NOT boil the agar.<br>Combine all ingredients. |
| 10 | Distribute the media to smaller bottles (if desired) and autoclave. |

Supplements should be added to agar medium while cool but still molten (approximately 50 °C), and the medium mixed thoroughly before being poured into plates.

**Supplements for leaf press plates include:**

- methanol (MeOH) [carbon substrate]: add 5 mL of 100% methanol per L of MP to reach a final concentration 125 mM
- cycloheximide [inhibits fungal growth]: purchase or make stock solution; add sufficient stock to reach a final concentration in medium of 50 μg/mL
- RPMI 1640 Vitamins Solution [facilitates growth of diverse organisms]: May be purchased commercially as 100x stock, sold as an ingredient for Roswell Park Memorial Institute (RPMI) 1640 medium. Add 10 mL of 100x stock per 1 L of medium.
- $LaCl_3$ (lanthanum chloride) [selective for some methylotrophs; used in only some plates]: make stock solution in $ddH_2O$; add sufficient stock to reach a final concentration in medium of 2 μM

#### Appendix 3. An abstract-style writeup assignment as a summative assessment for Methylothon, given in a Biotechnology class

### *Methylobacterium* Abstract

**Your completed written work cannot exceed 2 pages.**
**This does not include the phylogenetic tree or reference page.**

**Your final work must include the following areas:**

1. **Introduction:** An intro to *Methylobacterium* that addresses the following:
    a. What *Methylobacterium* are (identify characteristics that define the classification (see https://microbewiki.kenyon.edu/index.php/Methylobacterium for help).
    b. Major current work (research and/or application) involving *Methylobacterium*
    c. Purpose of our experiment
2. **Protocol/Methods:** A brief description of the experiment - include all major work involved in this lab, including stuff we did not directly do. Be brief but concise - a fellow scientist should know exactly what you did but you do not need to provide all the details.
    - **Note**: Although we did not do a lot of actual lab work, I would like you to describe the things we did cover, which will include the **sample collection and culturing** (leaf press and culturing after), **DNA extraction** (we did not fully cover this so very broadly what does DNA extraction mean here), **PCR** (which we did not do but you should be able to describe broadly how this should work, remember we would be looking to amplify certain sections of our microbes genomes to be able to compare them), and **sequencing** (which we did not do but you should be able to briefly explain what would happen in a sequencing reaction if you did complete it (not the lab part but how does sequencing work, broadly). **Just to recap -- you do not need to explain HOW these things would work in the lab but conceptually what is the purpose of each.**
3. **Results:** Attach a screenshot of your phylogenetic tree and then briefly describe your portion of the phylogenetic tree. This section is just you presenting what you found.
4. **Conclusions:** Provide a 2-3 sentence summary on what the results actually mean. **Address this idea in your conclusion:** Microbes are incredibly diverse and we know almost nothing about them. (How does your work relate to this statement and either support it or refute it?)
5. **Research Question**: Answer a question of your own that relates to the work we did. This should be related to the topics we covered in class but can be about anything of interest to you. This must involve outside research of primary or secondary sources. We will cover what this means in class. See this document for your opening question: Opening Questions (you can use one from here or a question of your choice). You should integrate your research into the work we did as best you can. Example: if your question deals with applications, then make sure to make the connection between applications and the work we did.)
6. **Citations/References**: Include a reference section that is set up using APA format.

| **Abstract Scoring Guide** | |
|---|---|
| **Introduction includes**<br>● A thorough but concise overview of *Methylobacterium*<br>● Discussion of current work/research and use of *Methylobacterium*<br>● The purpose of our experiment | **/20** |

| | |
|---|---|
| **Protocol includes**<br>● A basic explanation for each major section of the experiment (that does not get bogged down in details). | **/5** |
| **Results**<br>● Includes observations from the parts of the experiment we did (from the sampling and plate observations; you should include pictures.)<br>● Discusses and cites specific data (from the sampling, the plate observations and bioinformatics) | **/20** |
| **Conclusions**<br>● Summarizes what your tree tells you and its implications to microbial diversity | **/25** |
| **Research Question**<br>● Question is interesting and related to the topic<br>● Answer is thorough and uses primary and secondary sources to support it (you do not have to fully answer the question if it is something that is still an ongoing area of discovery) | **/25** |
| **References**<br>● **Only 1 verified reference is required.** | **/5** |
| **TOTAL:** | **/100** |

**Appendix 4. A Team Quiz as a summative assessment for Methylothon, assigned in an International Baccalaureate Biology class.**

#### QUIZ FOR THE METHYLOBACTERIUM WEEK

____/4 PROFICIENCY POINTS

From Dr. Martinez-Gomez, you have learned that some species of *Methylobacterium*, but not all, require rare earth elements to survive. We provided you with two culture plates designed for isolating methylotrophic bacteria; the plates were identical except that one contained the rare earth element lanthanum and the other did not. You used the plates to isolate microorganisms from a plant leaf. ***Your final assignment is to work with your team to complete a 1-2 page limit (double spaced with 1 inch margins) writeup with the following:***

1. Background information (considering citing Monday's reading here, Ceci's presentation, presentations from the week)

2. Describe a straightforward **research question/hypothesis** that can be tested through the experiment that you carried out. NOTE: Your team needs to coordinate this BEFORE you choose your leafs.

3. BRIEFLY summarize your **procedure**.

4. Describe your **results/observations** of microbial growth on your 4 culture plates at the end of the experiment. Were your observations sufficient to address your hypothesis?

5. If so, please explain your **conclusions** and the evidence supporting them. Consider mentioning **sources of error, limitations to the experiment, improvements, and/or an extension.**
If not, that's okay! It can be very difficult to conduct rigorous scientific research in the space of 1 week! In this case, please explain why your results were inconclusive, and what you think we would need to do to change the experiment (assuming we had more time, more resources, and the ability to work in a lab) in order to truly address the hypothesis you posed about *Methylobacterium* and rare earth elements. You may choose to discuss some of the following concepts:
- the methods we use to identify microbes
- what we know microbial growth requirements
- the importance of replication
- qualitative versus quantitative observations
- the differences between getting results that you don't expect, getting insufficient information from an experiment, and experiment totally failing

**Make a title and attach your one-two page response to this google classroom assignment. Please include the names of all team members.**

#### Rubric:

| | 4 | 3 | 2 | 1 |
|---|---|---|---|---|
| Introduction | Presents a clear summary of the aims of the study and its significance. Includes a straightforward **research question and hypothesis.** Briefly describes experimental design. Probably includes **one or more references to supporting sources*.** | Either lacks clarity or is missing one of the primary elements. | Weak or missing primary elements | No real introduction. |
| Materials and Methods | Gives the reader a clear picture of the **methods and materials** used. Does not use prescriptive language. Uses specific, not general, terminology. Detailed, step-by-step procedures are clearly referenced. Avoids long, redundant descriptions | Some methods are presented so briefly and/or vaguely that it is unclear how or why they were done. May be some written as a protocol rather than a description. | Some methods are omitted; others are presented in a piecemeal, vague form. | Methods barely mentioned. |
| Results | All **results** are clearly presented, with a logical sequence. Controls are clearly indicated (*if applicable*). | Some data may be missing. | Data is presented haphazardly. It is sometimes not possible to tell what material or procedure was used to obtain the data. | No logical connection between methods and data. Irrelevant data may be included, and relevant data left out. |
| Discussion | It is clear that the methods and results have been understood. The results (including controls) are related to the questions posed and **analyzed** for their | There may be some lack of clarity. Did the writer understand why certain methods were used, and how the | Very little analysis of the results. Statements are vague and general. Inconsistencies | Mostly a restatement of results. No analysis given. No recognition of error sources. |

| | | | | |
|---|---|---|---|---|
| | effectiveness. Scientific reasoning is included. **Possible explanations for inconsistencies and/or unexpected results are given.** | results could shed light on the questions asked? Incomplete analysis of inconsistencies and unexpected results. | are explained by 'human error' or something similar. | No understanding of controls. |
| Cohesiveness | It is clear that the report covers a group of related procedures with a clear set of goals. | Sometimes the goals are not clearly related to the report. Some fragmentation occurs, with methods and results apparently unrelated to each other. | Transitions are abrupt. Each day's work seems unrelated to the next's. Aims are not clearly present throughout. | Disjointed. No flow. Very little use of headings, or explanatory sentences. |
| Spelling and grammar | No spelling or grammatical errors | An occasional error. | Apparently not proofread for errors. | Frequent grammatical errors: incomplete sentences, tense changes, misspellings. |

*Please cite any sources in APA. For example, if you use the Monday reading about lanthanides, please cite it in APA at the end.

**Appendix 5. A flexible-format final assignment combining Methylothon with human ancestry, given as a summative assessment in a Biotechnology class.**

#### Paleogenetics Project

You are assisting a team of anthropologists studying an ancient cave site. Nearby, burial grounds are discovered! These bones look different from known hominins (human species) in the area. A sample of bone containing DNA is given to you to analyze.

Because you are the team's expert in bioinformatics, it is your job to determine whether this sample comes from a known species of humans or an entirely undiscovered species. You'll also determine which hominins this group is most related to (where this group resides on the tree of life) and this group's likely migratory pattern (how these people came to this cave site).

Create a presentation* in which you answer these questions:

1. How will you obtain enough DNA for the analysis?
2. How will you prevent contamination?
3. How will you sequence the DNA?
4. How will you determine if this is a known species or a new species?
5. How is a multiple sequence alignment done, and how is a phylogenetic tree created?
6. How will you determine this group's migration route?

*You may choose to make a video, a slideshow, write an essay, or create a graphic novel. If you have another idea, run it by me, and we'll see if it could work.

Standards which apply to this assignment:

- I can describe the use of common lab equipment and sterile techniques.
- I can describe how DNA is isolated from a sample.
- I can describe the steps of PCR.
- I can explain how DNA is sequenced.
- I can explain what it means to BLAST a DNA sequence.
- I can describe what a computer does to create a multiple sequence alignment.
- I can create a phylogenetic tree using DNA sequences.
- I can describe what a molecular clock is and how it is used.
- I can explain how we determine human ancestry and migration.
- I can describe why mtDNA is a useful tool for studying ancestry.
- I communicate clearly and concisely about science.

#### Paleogenetics Project Rubric

| | **Exemplary** | **Proficient** | **Developing** | **Support Needed** |
|---|---|---|---|---|
| ***I can describe the use of common lab equipment and sterile techniques.*** | Excellent descriptions of lab equipment and sterile techniques. | Correct descriptions of lab equipment and sterile techniques. There are a few errors or omissions. | Incomplete descriptions: just lab equipment or just sterile techniques. | Incorrect terms or descriptions of equipment or sterile techniques. |
| ***I can describe how DNA is isolated from a sample.*** | Accurate and complete description of DNA extraction. | Generally correct description of DNA extraction. There are a few errors or omissions. | Incomplete description of DNA extraction.. | Incorrect description of DNA extraction. |
| ***I can describe the steps of PCR.*** | Accurate and complete description of PCR. | Generally correct description of PCR. There are a few errors or omissions. | Incomplete description of PCR. | Incorrect description of PCR. |
| ***I can explain how DNA is sequenced.*** | Accurate and complete description of DNA sequencing. | Generally correct description of DNA sequencing. There are a few errors or omissions. | Incomplete description of DNA sequencing. | Incorrect description of DNA sequencing. |
| ***I can explain what it means to BLAST a DNA sequence to create a multiple sequence alignment and a phylogenetic tree.*** | Accurate and complete description of DNA databases, the BLAST tool, MSA, and tree creation. | Generally correct description of DNA databases, the BLAST tool, MSA, and tree creation. There are a few errors or omissions. | Correctly describes two of the four. | Mentions DNA databases but not how they are used. |
| ***I can describe what a molecular clock is and how it is used to study human ancestry and migration.*** | Accurately explains how mtDNA functions as a molecular clock and how haplogroups are used to study human ancestry and migration. | Correctly describes how haplogroups are used to study human ancestry and migration. | Correct description of human ancestry or human migration. | Mentions mtDNA or haplogroups, but does not describe what they are or how they are used. |
| ***I communicate clearly and concisely about science.*** | The work flows, is clear and concise, and articulately explains each element. | The work is clear and concise. A few incorrect terms or grammatical problems. | The presentation gets off topic. Distracting errors in terms and grammar. | The work is muddled and contains many errors. |
| ***I turn in my work on time.*** | Assignment is turned in early | Meets deadline | Within 2 hours after deadline | Turned in late |

**Appendix 6. Plants sampled by students for leaf presses during Methylothon 2021.**

| |
|---|
| Abutilon sp. (Flowering Maple) |
| Abutilon sp. (Mallow) |
| Acacia melanoxylon (Black Acacia) |
| Acer palmatum (Green-leaf Japanese Maple) |
| Allamanda blanchetii (Purple Allamanda) |
| Anthurium andraeanum (Flamingo Flower) |
| Artemisia vulgaris (Common Mugwort) |
| Camiella japonica (Japanese Camellia) |
| Chrysanthemum sp. |
| Citrus limon (Lemon) 'Meyer,' 'Eureka' |
| Citrus x sinensis (Orange) |
| Claytonia perfoliata (Miner's lettuce) |
| Crassula multicava |
| Cyclamen persicum (Persian cyclamen) |
| Delairea odorata (Cape Ivy) |
| Fern, unknown species |
| Ficus benjamina (Weeping Fig) |
| Fortunella japonica (Kumquat) |
| Fragaria sp. (Strawberry) |
| Fragaria vesca (Wood strawberry) |
| Gaultheria shallon (Salal) |
| Geranium purpureum (Little-Robin) |
| Geranium sp. |
| Ginko biloba |
| Hedera helix (common English Ivy) |
| Hibiscus sp. |
| Hoya carnosa (Honey plant) |
| Impatiens sodenii (Poor Man's Rhododendron) |
| Ipomoea purpurea (Common morning glory) |
| Lilaceae sp. (Lily) |
| Lonicera japonica (Honeysuckle) |
| Loropetalum chinense (Fringe flower) |

| |
|---|
| Macadamia sp. |
| Medicago lupulina L, (Black Medick) |
| Mentha (Mint Leaf) |
| Nepenthes x Miranda (Tropical pitcher plant) |
| Oxalis pes-caprae, (sour grass, Bermuda buttercup) |
| Oxalis stricta (common yellow woodsorrel) |
| Pachira aquatica (Provision Tree) |
| Parietaria sp. |
| Pelargonium peltatum (Ivy Geranium) |
| Persea americana (Winter Mexican Avocado) |
| Persicaria sp. (Knotweed, Smartweed) |
| Philodendron hederaceum (Heartleaf Philodendron) |
| Philodendron laciniatum |
| Physalis peruviana (Cape Gooseberry) |
| Pisum sativum (Common Pea) |
| Pleargonium zonale (L.) L'Her. ex Aiton " (Horseshoe geranium) |
| Quercus agrifolia (Coast Live Oak) |
| Quercus sp. (Oak) |
| Rosa sp. (Rose) |
| Rubus Armeniacus (Bramble, Himalayan Blackberry) |
| Spathiphyllum sp. (Peace Lily) |
| Tagetes erecta (Orange Marigold) |
| Trifolium sp. (Clover) |
| Tropaeolum sp. (Nasturtium) |
| Urtica urens (Dwarf Nettle) |
| Uvularia grandiflora (Yellow Bellflower) |
| Vicia faba (Fava bean) |

**Appendix 7. Example of student work on Bacterial Identification Virtual Lab worksheet.**

**Methylothon**
**Bacterial Identification Virtual Lab** **Date: ____10 March 2021____Period: ___**

**_____/4 POL Points**

| **Team Member Names and Roles:**<br>[redacted] |
|---|

First, **document managers, make a copy** of this document and share it with your group so that all of you can add to the document at the same time.

Next put your name in the blank:

______ **scientist:** shares screen and manipulates lab (you must have Flash)
________ **recorder:** writes down the procedure (tells the story)
______ **documents manager:** screen captures pictures of the tools, results
_______ **captain:** writes down what the tools are used to do

Scientist: access the virtual **Bacterial Identification Lab** and share your screen. Follow the instructions and click the prompts.

Recorder, documents manager, and captain: split your screens so that you can watch the lab AND edit this document at the same time.

***A. Sample Preparation***

1. --Documents manager: **upload** a picture of the sample plate with bacterial colonies.

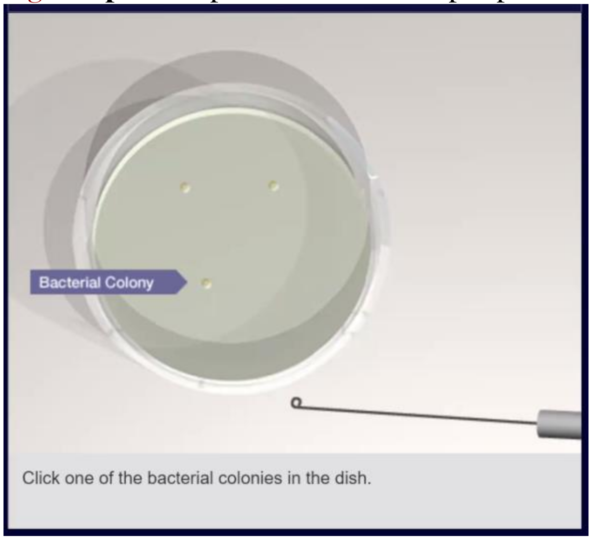

2. --Captain: What is the purpose of centrifuging the sample? Which part of the centrifuged product are we collecting?

| |
|---|
| We centrifuge to separate the sample by weight in order to get rid of the proteolytic enzyme. By centrifuging the sample, the cellular debris is removed from the sample. We are collecting the DNA liquid that is left in the centrifuge. |

Everyone:

3. --In Methylothon, we isolate our *Methylobacterium* with a specific kind of culture plate. What kind of medium do we use, and why?

| |
|---|
| We use a selective minimal culture medium in order to only allow certain organisms to grow. In the case of the lab, our specific medium is designed to allow for only the methylobacterium to grow. |

4. --Why is it important to select a sample from one colony, rather than collecting from multiple colonies?

| |
|---|
| Different colonies could have different methylobacterium on them and we only want to see the methylobacterium from one colony, not many. |

***B. PCR Amplification***

5. --Recorder: list the times and temperatures for each step of the PCR process

| |
|---|
| Denaturing-Temperature: 95°C, Time: 5 minutes on initial cycle.<br>Annealing-Temperature: 5°C below Tm of primers; no lower than 40°C. Time: 30-45 seconds.<br>Extension- Temperature 72C, Time: ~1 min/kb of expected product; 5-10 min on last cycle. |

6.--Captain: List the substance in the red, green, and blue vials and explain why they are used in the experiment.

| |
|---|
| Red: the red is the PCR master mix and is used to keep the pH constant<br>Green: the green is the positive control DNA, and is used as a control<br>Blue: the blue is deionized water, and it is used as a negative control |

7.--Documents manager: capture images of the PCR animation

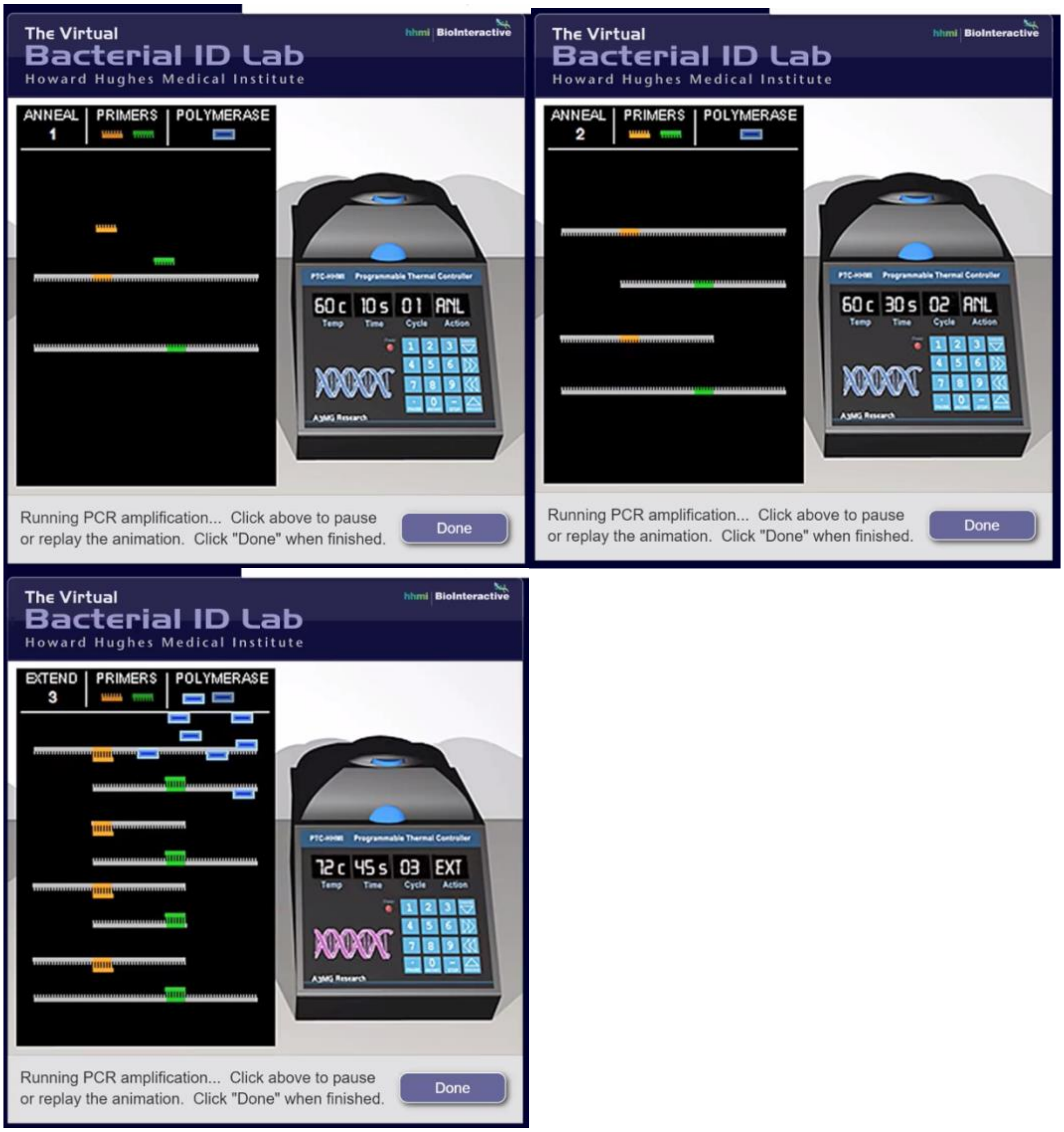

Everyone:

8. --Explain the significance of the temperature required by each step.

Denaturation – Temperature: 95°C  is used to separate the double helix DNA strands
Annealing – 60°C is used so the primer can bind to the single stranded DNA (annealing)
Extension – Temperature 72C is used for the creation of new strands of DNA made using the original stands as templates.

9.--What is the purpose of using heat-stable DNA polymerase in PCR reactions?

Since 95°C is the denaturing temperature, and 5°C below Tm of primers is annealing, the polymerase must be able to not denature in these temperatures due to the role it plays in separating, amplification, and reconstruction.

10. --How many copies of DNA are generated at the end of 30 cycles?

There are 1 million copies at the end of 30 cycles.

***C. PCR Purification***

11. --Recorder: List the steps in PCR Purification

1. PCR Buffer PB (400ul)
2. Column binding
3. Washing
4. Drying
5. Elution
6. Pure DNA fragment

12. --Captain: What additional substance are we adding to the column? Why?

We are adding a buffer solution to the column because it makes sure that the DNA can separate from the column and go into the collection tube.

13. --Documents manager: take an image of the column after it is transferred to a new test test tube. Which tube contains the supernatant? What is in the supernatant?

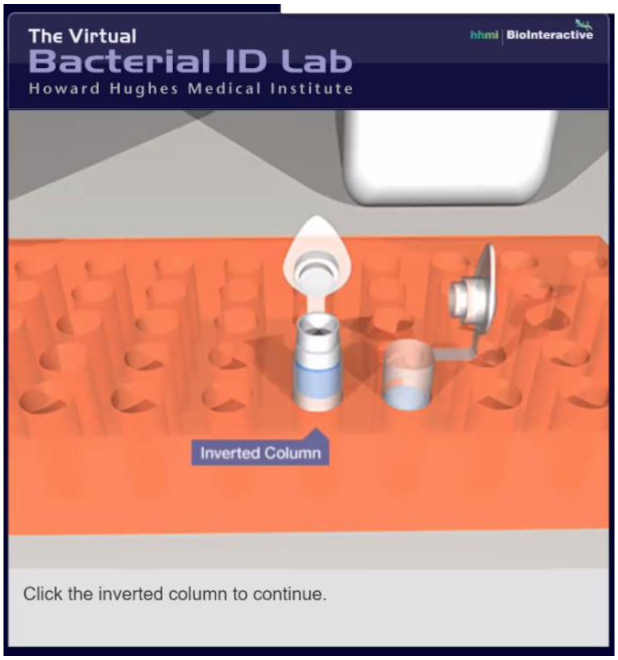

The old test tube would contain the supernatant since the column was removed and placed into a new test tube, separating the DNA and leaving the supernatant at the bottom of the old test tube. The supernatant is everything but the DNA.

Everyone:

14. --Explain the results we expect to see in each of the three lanes of our gel electrophoresis and why these expected results would indicate a successful PCR reaction.

One lane would be a negative control only containing water.
The middle lane would be a positive control containing the PCR known product.
The last lane would be for the unknown sample.

The results that would indicate a successful PCR reaction would include a positive control reaction.

15. --What alternative to gel are we using to purify our product?

Compact microfilters is an alternative to gel.

***D. Sequencing Prep***

16. --Captain: Explain what is in the "Sequencing brew" in the blue and green tubes.

The green and blue tubes had different buffers and primers in each tube in addition to DNA polymerase and fluorescence tagged terminators.

17. --Documents manager: Take a picture of the different length sequences. **Paste** it below.

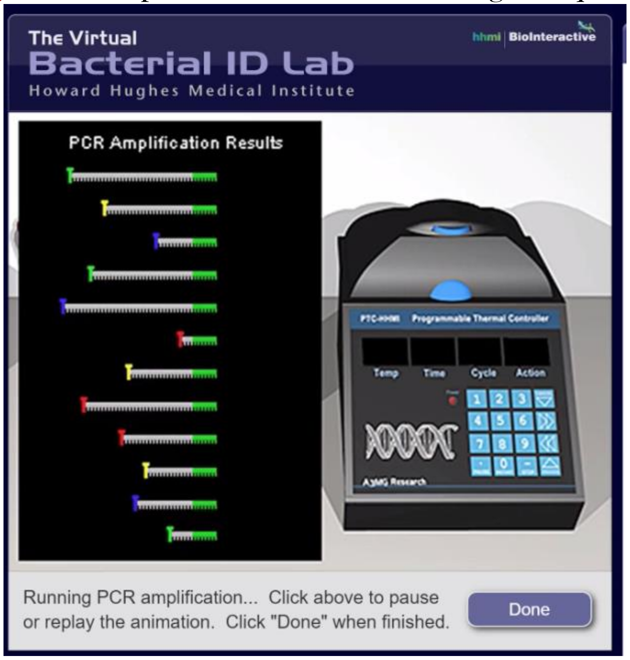

Everyone:

18. --Why do we use multiple primers in PCR cycle sequencing for long sequences?

Using multiple primers allows many short, overlapping pieces of DNA to be put together to find the complete sequence.

19. --What is the significance of using primers that bind to conserved regions of the 16s rDNA gene?

This allows them to bind to the sequence regardless of bacterial source.

20. --Describe what occurs in the tube containing the primer 651R.

In this tube, the DNA strands bind to the primer and have one fluorescence-tagged terminator at the end that they don't bind to the primer.

***E. DNA Sequencing***

21. --Documents manager: capture a picture of the chromatogram. **Paste** it below.

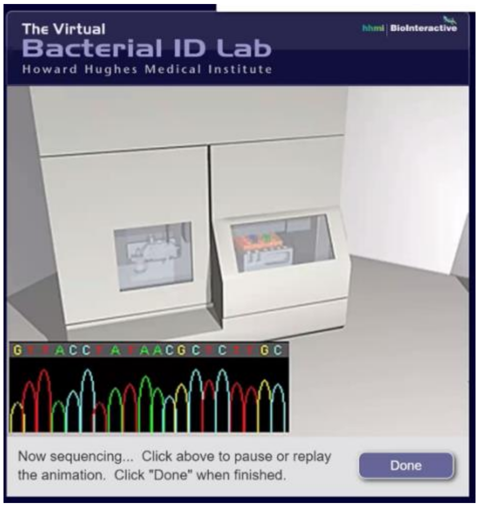

Everyone:

22. --Explain why DNA molecules move from one end of the gel to the other during gel electrophoresis.

| |
|---|
| Gel electrophoresis applies an electrical current to the tube, and since DNA molecules are negatively charged, they will move through the tube towards the positively charged end, smaller pieces moving faster. |

23. --What is the purpose of fluorescent markers in DNA sequencing?
In what order do DNA fragments move through the sequences (i.e. why might one sequence travel faster than another)?

| |
|---|
| The fluorescent markers are used when the DNA fragments are pushed through a laser beam and this interaction sparksdecistors to recognize the fluorescent markets. The DNA fragments are pushed through the gel based on size. |

***F. Sequencing Analysis***

24. --What species of bacteria does the DNA belong to?

| |
|---|
| The DNA belongs to the Bartonella henselae species. |

Documents managers, please turn in this assignment on google classroom. Everyone else, please mark this as done WITHOUT turning it in.

**Appendix 8. Example of student work in Bioinformatics Virtual Lab worksheet.**

Name: Date: 3-12-21 Period:

### *Bioinformatics Tutorial Worksheet (Thursday Class and Lab)*

_____/4 POL Points

*In this BLAST tutorial, we're going to be downloading files, uploading files, and moving from website to website frequently. There's a lot of moving parts here, so this worksheet (hopefully) will help make some of the nitty gritty website-wrangling steps clearer. Of course, feel free to stop by office hours if you have any questions!*

#### *1. Getting your sequences for BLASTing*

In this lab, you'll be BLASTing an unknown sequence collected from a plant leaf by last year's Methylothon-ers! You'll also need reference sequences for building a phylogenetic tree in the final step.

1. Navigate to http://methylothon.com. Find your individual assigned DNA sequence here.
2. From the main menu, go to "Methylothon 2021: student page"
3. Sign in with the password PinkBacteria2021
4. Follow the instructions on the website under "DNA sequences you need for the Bioinformatics Tutorial".

#### *2. BLASTing your unknown*

*What's happening in this step:* In this step, we're using NCBI's BLAST tool to compare our unknown DNA to a database of known sequences. We're going to gather the top hits (known sequences in the database most similar to the unknown sequence) and construct an alignment.

first, navigate to the BLAST homepage at https://blast.ncbi.nlm.nih.gov/Blast.cgi

scroll down and click the "Nucleotide BLAST" button.

your screen should look like this:
click the "choose file" button (circled in red) and upload your unknown.

scroll down, and make sure that the "megablast" option is enabled.
then, click BLAST to see your results!

****sometimes, BLAST will take a while to search and you'll get stuck on this loading screen for a minute -- this is normal!****

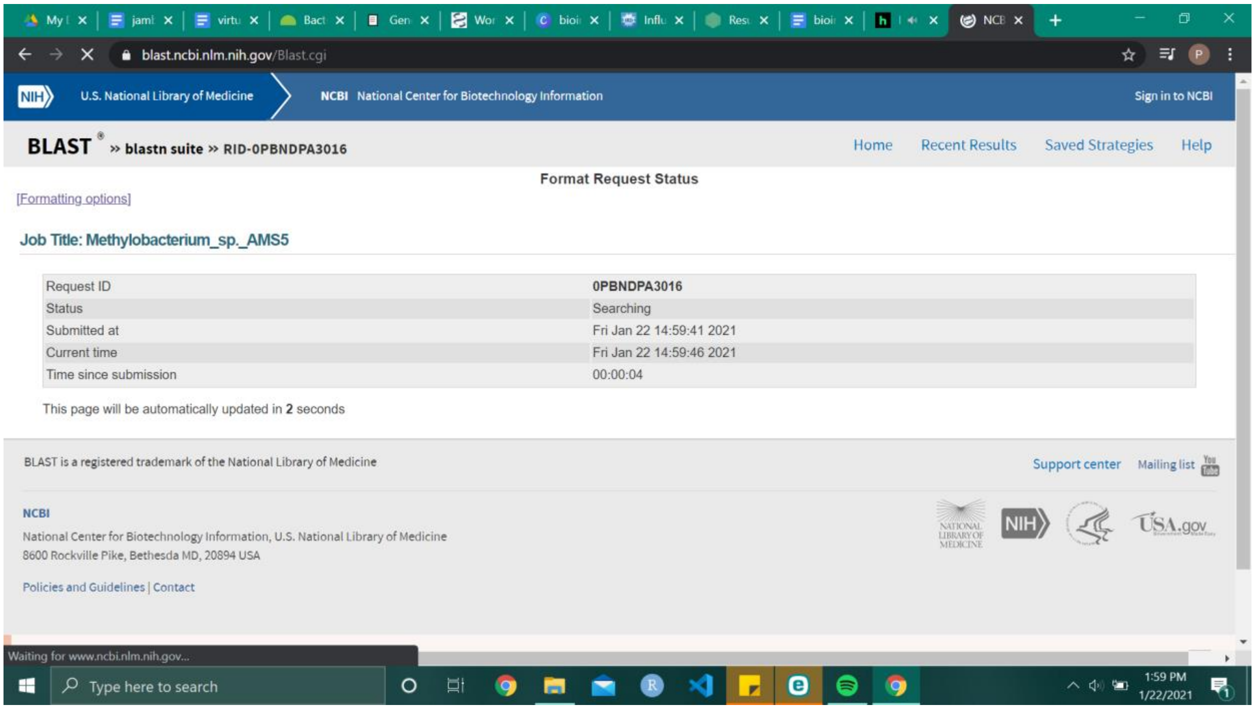

scroll down on the results page until you see a list of match results:

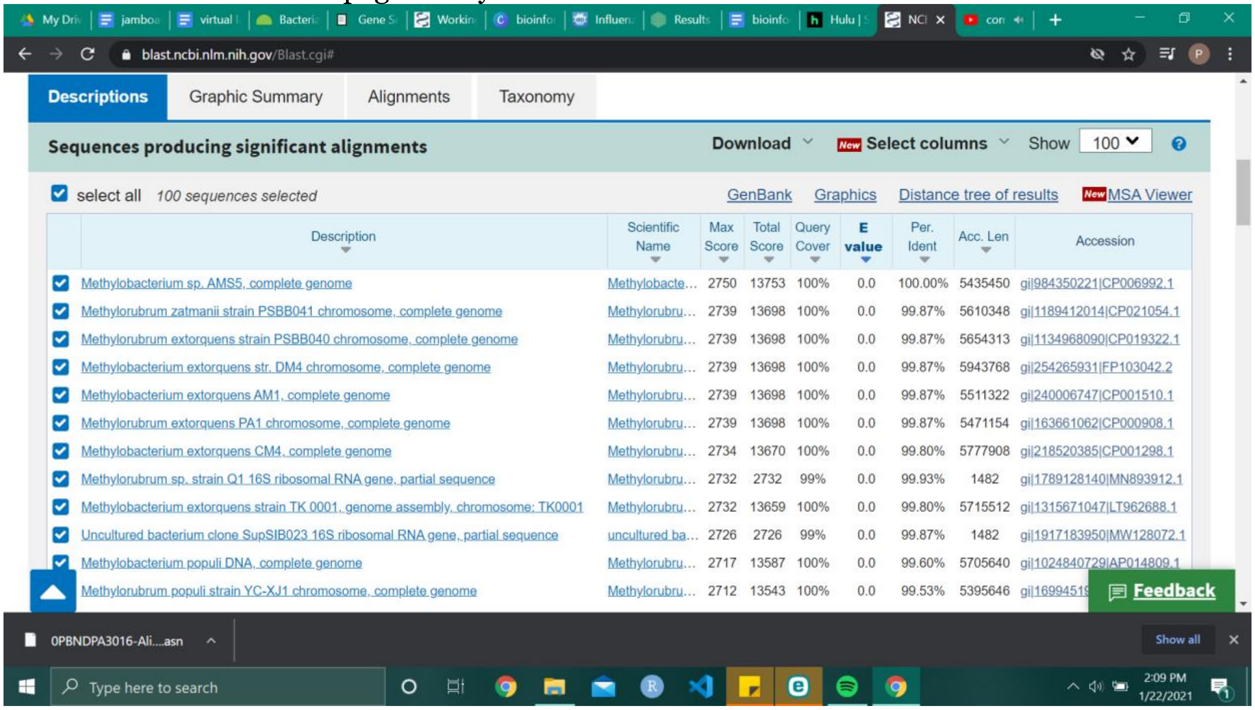

uncheck the "select all" button, and manually check the first 4 results.

click "download" and download the *aligned* FASTA sequences. We choose this option so that we download the only portions of the sequences that match our query sequence in length.

***A. STOP AND ANSWER: What kinds of organisms are the top hits for your sequence? Is this what you expected? Explain. If you're curious, you can click on any of those hits and follow the weblinks to the Nucleotide database entry. This entry sometimes contains information about where the sequence came from.***

| |
|---|
| The organisms that were top hits for the sequence of my DNA sample were all sequences of the ribosomal RNA gene of methylobacterium. This is what I expected because the lab that was done in order to obtain this DNA was one that was intentionally designed to culture methylobacterium. The sample this student collected from their plates contains DNA of the methylobacterium that this student cultured from their leaf press. |

***2.5 Reformatting***

Before we visualize relatedness among our three different DNA sources (your original unknown, top BLAST hits, and reference sequences), we can make the process of managing computer programs much easier if we condense all of our sources into a single file. The files we'll be bundling together are all FASTA files. A FASTA file is a filetype that displays species name / information followed by genetic code. An example of the FASTA format is provided below. FASTA files are often saved with the file extension ".fasta" or ".fa" to help DNA-sequence-editing programs recognize what they are, but they can also be saved with the extension ".txt", which is what we will do here.

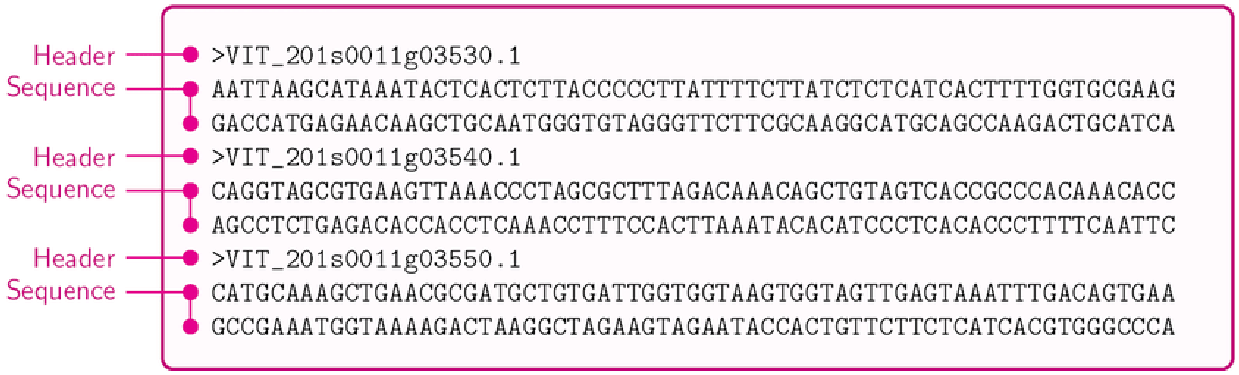

We're going to create a file that has all the DNA we want to investigate in a single file. To do this, open up a plain text editor (for instance, "Notepad" on Windows devices, or "TextEdit" on Mac). Then open up the following:

1. your BLAST top hits (called "seqdump.txt" unless you renamed it)

2. the reference sequences you downloaded from the project website (called "Methylothon_2018-2019_reference_sequences.txt")
3. your mystery sequence.

Paste in all of your sequences of interest from the three different files into a new document, one right after another as seen in the example image above (eg. no blank lines between sequences).

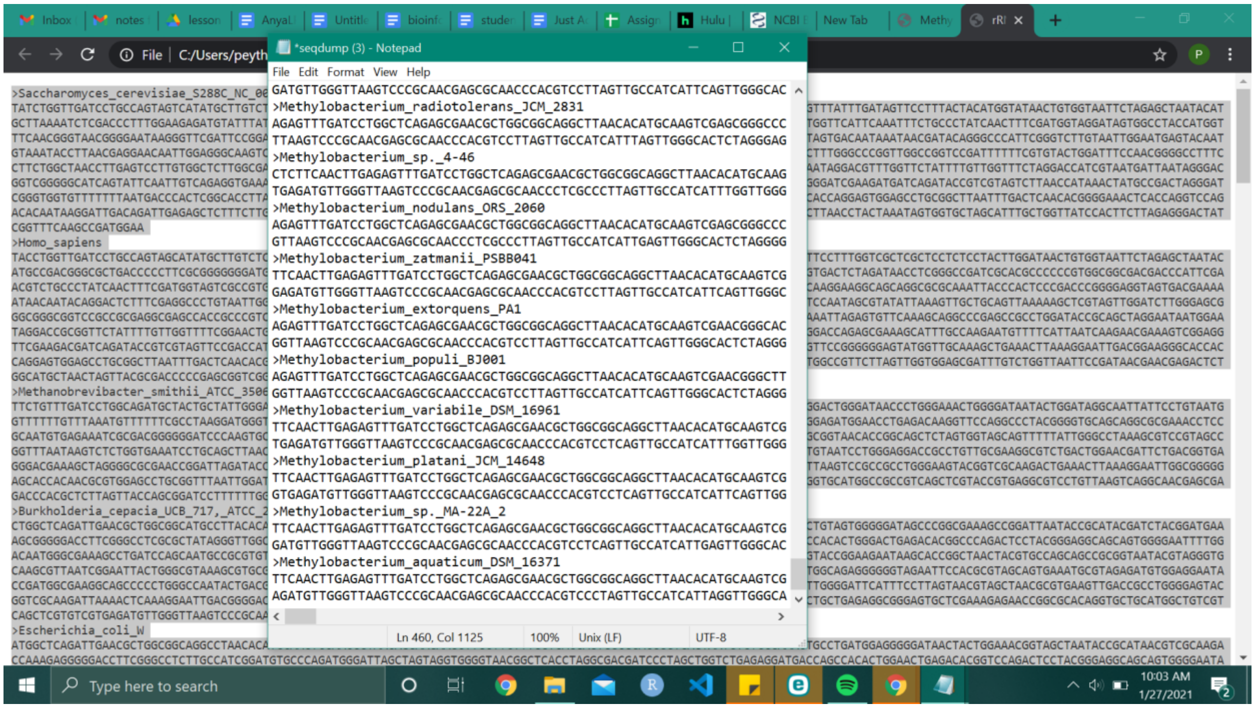

Save your new document as [LASTNAME_CondensedSeqs].txt.

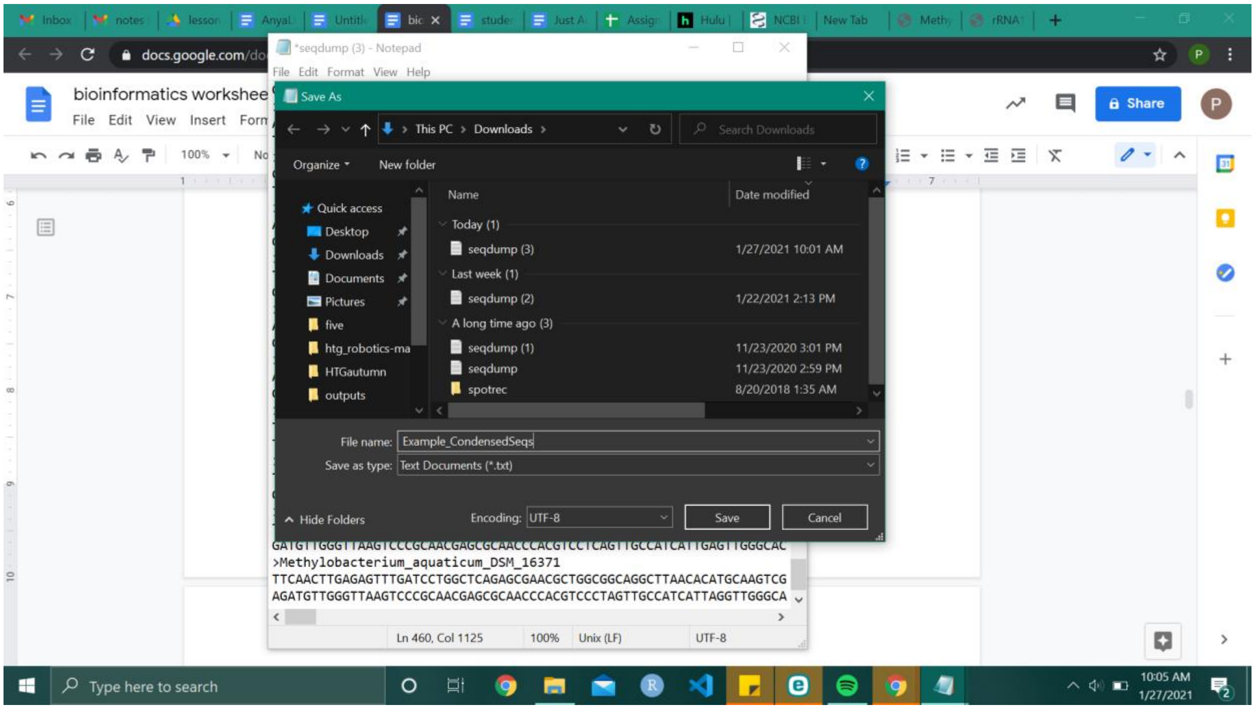

##### 3. *Multiple Sequence Alignment*

*What's happening in this step:* In this step, we're going to visualize the alignments of our unknown and the closest hits, as well as our reference sequences.

We're going to use EMBL's Clustal Omega tool at this link:
https://www.ebi.ac.uk/Tools/msa/clustalo/

First things first, we're going to change the set from "PROTEIN" to "DNA"
upload your combined FASTA file, scroll down and click submit!

to view a MSA of your results, click the "results viewer" tab.

Scroll down to "View in MView" and click it!

the MSA generator will be autopopulated with the data from previous step...

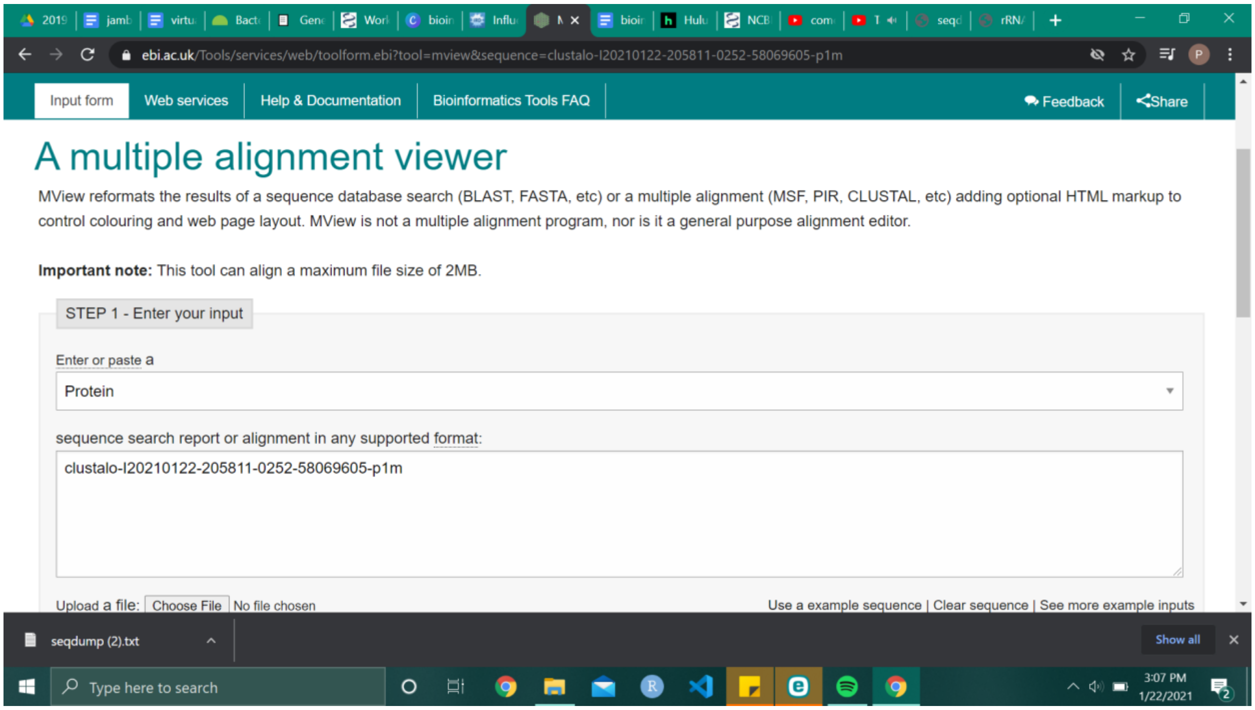

so just scroll down and click submit!

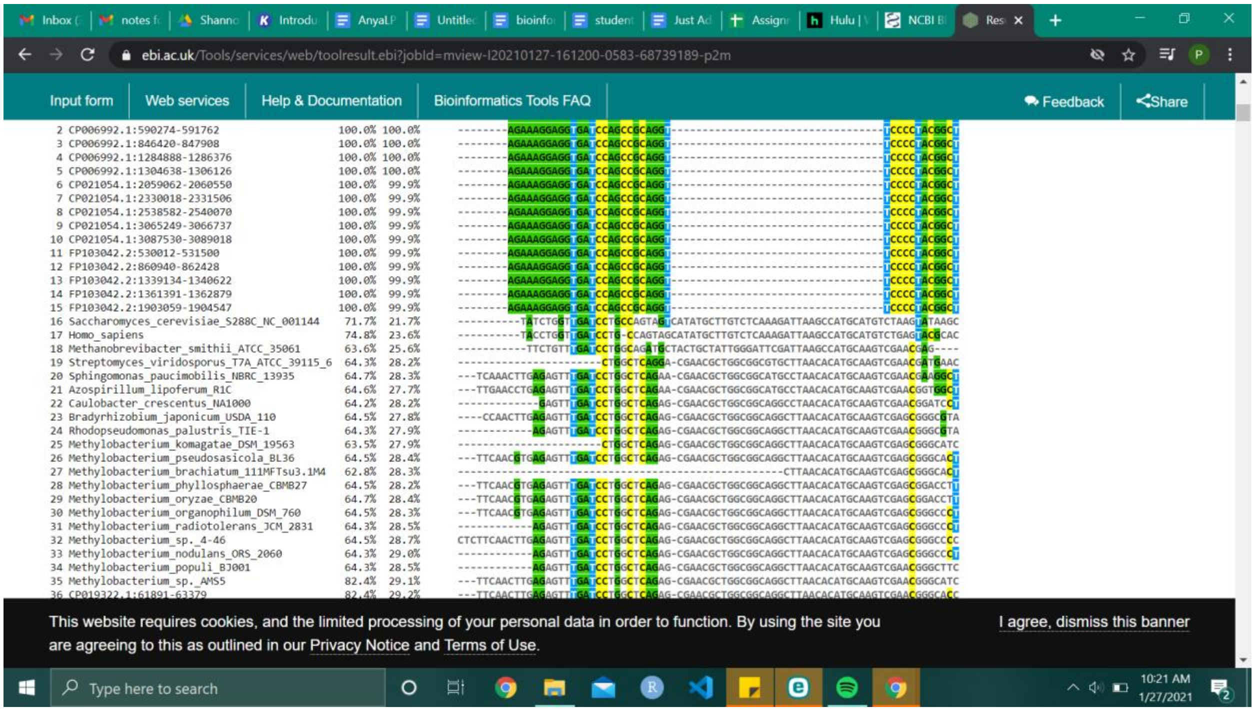

***B. STOP AND ANSWER: Inspect the alignment.***
***Do the sequences generally look pretty well aligned? How can you tell?***

| |
|---|
| Yes, the sequences look well aligned, made evident by the highlighted columns of nucleotides. This is especially true for certain sets of samples compared to others, as some groups have less alignment than others. |

C. ***STOP AND ANSWER**: Just from looking at the alignment, can you guess which sequences are most distantly related from your mystery sequence?*

| |
|---|
| There are certain methylobacterium samples that align less than others, and these sequences, along with the sequences pertaining to species that are not methylobacterium, are most distantly related from my mystery sequence. |

#### *4. Phylogenetic Tree*

*What's happening in this step*: Now, we're going to create a visualization of relatedness among the top hits and the reference sequences.
To do this, we'll be using the Influenza Research Database's tool found at this link:
https://www.fludb.org/brc/tree.spg?method=ShowCleanInputPage&decorator=influenza

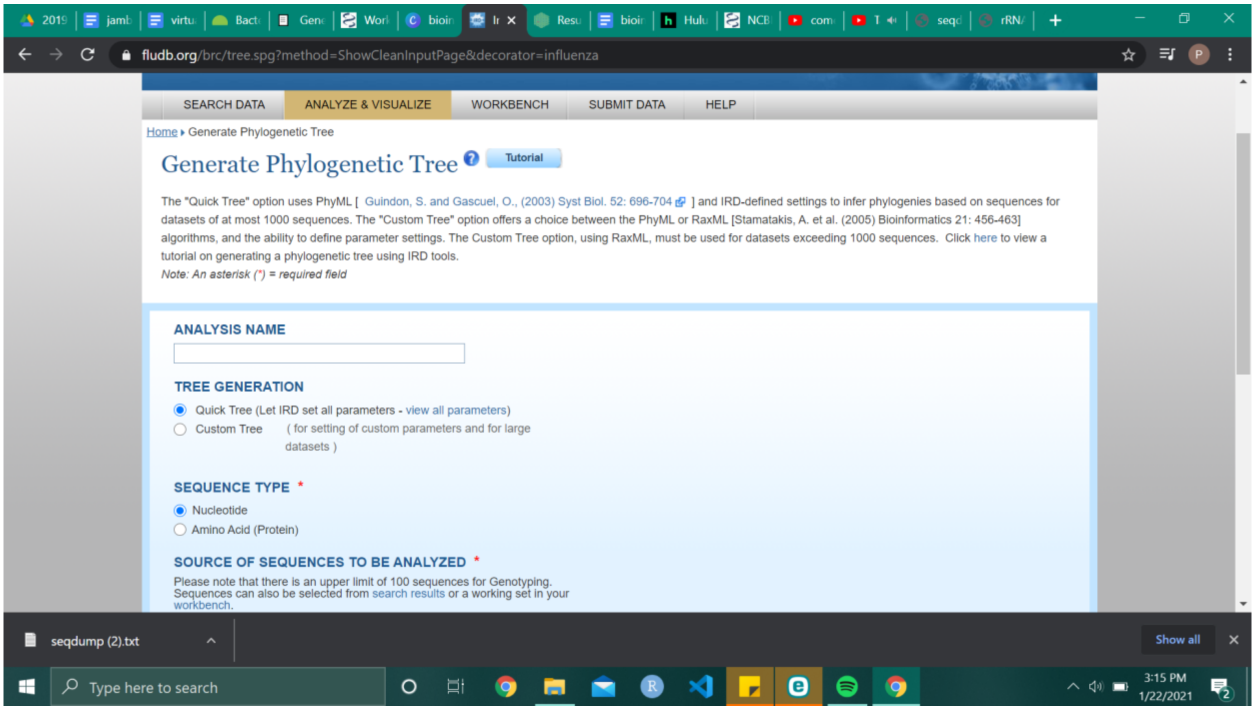

same as before, we'll be uploading our combined FASTA. make sure that you check <u>unaligned</u> FASTA format, and click submit!

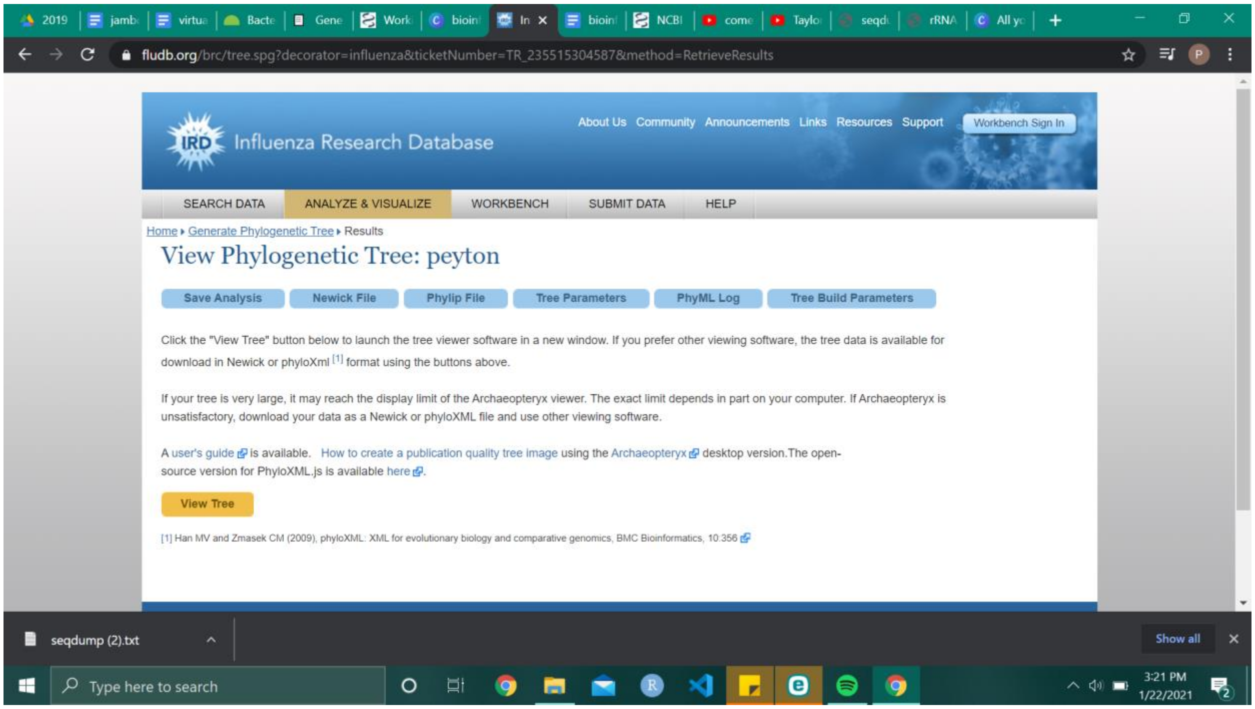

click view tree, and check out your phylogeny!

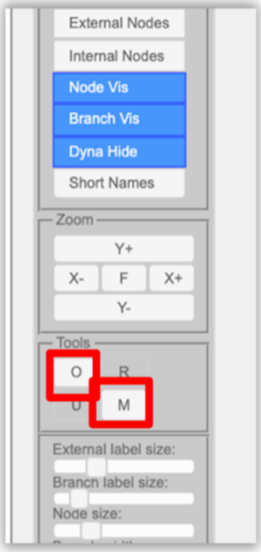

There are two quick manipulations you should carry out to make the tree easier to compare with others in your class.
First click the "M" button for "mid-point re-root" - this makes a guess at where to place the ancestor.
Then click the "O" button for "order all nodes" - this prettifies things by ordering branches by length (but doesn't change the interpretation of the tree).

Enter the name of your top blast sequence in the search bar to highlight it, and download an image of your tree with your highlighted sequence.
then you're ready to upload the image to the class slideshow!

***SUBMIT YOUR TREE: Follow this link to the class slideshow. Find the slide with your name on it and import the image of your phylogenetic tree. Add the information about your isolate's host plant.***

D. ***STOP AND ANSWER: According to the tree, what are your mystery organism's closest relatives? Is this what you expected? Explain.***

My mystery organism's closest relatives are other sequences of the ribosomal RNA gene of methylobacterium, according to BLAST results. According to other reference species on my tree, the other closest relatives to my isolate are other varying species of methylobacterium including *brachiatum, pseudosassicola*, and *phyllosphaerae. Yes, seen as my isolate is a methylobacterium, it was expected that its closest relatives would also be.*

E. ***STOP AND ANSWER: Once student results begin to fill the powerpoint, answer the following questions:***

a) Find another student who has an isolate that appears genetically identical to yours.
What is the name of that isolate? 2-ZX
Compare sequences-- just give them an initial glance. Do they look like they might be the same? Explain.

No, these sequences aren't exactly the same, but they are very similar in terms of alignment. Many nucleotide bases align with each other, with a few exceptions of substitutions and gaps. The differences increase towards the end of the sequence.

If you really wanted to know for sure whether they were exactly the same all the way through, how would you do that?

In order to do this, I would simply repeat this process with the sequence of my isolate and this similar isolate. I would combine the sequences into a file and run it through the website that highlights corresponding pairs and similarities. The more highlighted these two sequences are, the more similar, and if they are completely highlighted, that means they are exactly the same all the way through.

b) Find another student who has an isolate that is not identical, but looks from your trees to be closely related.
What is the name of that isolate? 6-CL
Describe how you know that your sequences are closely related but not identical. You can base this claim purely on the phylogenetic tree, or also on the DNA sequence or BLAST results.

I know this sequence is closely identical to mine solely based on the phylogenetic tree. This isolate's closest relatives are also partial ribosomal RNA gene sequences, which are the closest relatives to my isolate as well according to BLAST results. Furthermore, at a node only one further back on 6-CL's phylogenetic tree, it branches off to other species of methylobacterium from the reference list, which are also close relatives to my isolate. This includes methylobacterium *brachiatum, pseudosassicola*, and *phyllosphaerae*.

c) Find a student who has an isolate that is not *Methylobacterium*
If your own isolate was not *Methylobacterium*, find a different student!
What is the name of that isolate? 4-MC
What kind of organism do you think that isolate is? Sphingomonas bacterium

d) Find a student who has an isolate from the same plant species as yours. [Note: not all plants had two isolates, so if you don't find a match, skip this question.]
What is the name of that isolate? 1B-CC
Are your isolates genetically similar? Explain how you know.

Yes, our isolates are genetically similar. Both our isolates are from the same plant species, which means the cultured organism is likely to be the same. This is confirmed by our closest relatives, which according to BLAST are the same. Not only this, but the references are organized in the same locations, with other species of methylobacterium similarly related to our isolates.

e) BONUS: Did anyone in your class find something that could potentially be a novel species of *Methylobacterium*? (Assume that the reference sequences we provided include all the known species of *Methylobacterium*, which is quite an assumption.) Explain how you would recognize that.

The isolate 6A-RM is its own outgroup when compared to methylobacterium, despite still being placed closely on the phylogenetic tree. This could possibly signify a novel species of Methylobacterium having been discovered.
